# The Anticipation of Events in Time

**DOI:** 10.1101/608893

**Authors:** Matthias Grabenhorst, Georgios Michalareas, Laurence T. Maloney, David Poeppel

## Abstract

Humans use sensory input to anticipate events. The brain’s capacity to predict cues in time is commonly assumed to be modulated by two uncertainty parameters, the hazard rate (HR) of event probability and the uncertainty in time estimation, which increases with elapsed time. We investigate both assumptions by manipulating event probability density functions (PDF) in three sensory modalities. First we show, contrary to expectation, that perceptual systems use the reciprocal PDF – and not the HR – to model event probability density. Next we demonstrate that temporal uncertainty does not necessarily grow with elapsed time but also diminishes, depending on the event PDF. Finally we show that reaction time (RT) distributions comprise modality-specific and modality-independent components, the latter likely reflecting similarity in processing of probability density across sensory modalities. The results are consistent across vision, audition, and somatosensation, indicating that probability density is more fundamental than hazard rate in terms of the neural operations determining event anticipation and temporal uncertainty. Previous research identified neuronal activitity related to event probability in multiple levels of the cortical hierarchy such as early and higher sensory (V1, V4), association (LIP), motor and other areas. This work proposed that the elementary neuronal computation in estimation of probability across time is the HR. In contrast, our results suggest that the neurobiological implementation of probability estimation is based on a different, much simpler and more stable computation than HR: the reciprocal PDF of events in time.

## Introduction

Perceptual systems – auditory, visual, or somatosensory – anticipate events. Successful anticipation at the second scale allows the organism to prepare a response before an event occurs. At one extreme, prediction of events may fail entirely: events arrive unexpectedly. At the other, sensory information serves only to confirm what was in effect already known based on accurate prediction, for example in the context of complete stimulus regularity. In between these extremes, information about the likely time of arrival of the next event can be summarized as a *probability density function* (PDF) across time. For the brain, the estimation of event occurrence is influenced by two main sources of uncertainty: first, by the actual probability distribution of events and second, by the brain’s inherent uncertainty in estimating elapsed time (1, 2). In previous work, ranging from single-cell recordings (3-7) to non-invasive electrophysiology (8, 9) and neuroimaging (10, 11), systems neuroscience has provided two compelling hypotheses for the computations involved in the neural representation of both sources of uncertainty.

*Hypothesis A* states that the brain models the probability distribution of event occurrence by computing the *hazard rate* (HR) (3-10, 12, 13), an intuitively pleasing and conceptually straightforward model of anticipation: the HR is defined as the probability density of an event at any point in time, given that it has not occurred before (14). Although the HR appears to represent precisely the information needed in temporal anticipation, one should be cautious in asserting its validity based on this fact alone (15). Technically, the hazard rate relates a PDF to its integral, the *cumulative distribution function* (CDF). More specifically, the HR is the PDF divided by the survival function: HR = PDF/(1-CDF) (14). Alternatively, the HR can also be interpreted as the PDF multiplied by 1/(1-CDF), rendering the reciprocal survival function a time-varying scaling factor for the PDF. This suggests that the fundamental variable in the HR’s equation is the PDF. Therefore, although the HR and the PDF are typically thought of as separate representations of probability density, they could be conceived as related variables, each possibly important in the brain’s effort to model event probability. *Hypothesis A* further posits an inverse relationship between reaction time (RT) and event expectation: The RT to an event is linearly anti-correlated with HR (5, 9) or, put differently, linearly correlated with the *mirrored* HR (reflected around a constant value). One significant challenge for *Hypothesis A* is that the computation of HR, which requires integration of event probability over time, is relatively complex and sensitive to noise (14, 16).

*Hypothesis B* assumes that the brain’s uncertainty in estimating elapsed time is increasing monotonically with time, which is often referred to as the scalar property of time estimation (17, 18). At the behavioral level (5, 6, 13) and at the neural level (3, 5-7, 19), this uncertainty is typically modeled as a Gaussian function whose variance increases with elapsed time, a computation which is termed here as *temporal blurring*. Critically, temporal blurring ignores potential effects of environmental temporal statistics on time estimation, which raises the question whether the brain’s estimation of time is influenced directly by event probability. To build on and extend the above concepts, we test probabilistic inference across time at the behavioral level (20). Guided by *Hypotheses A* and *B*, we examined the potential of several explanatory variables for modeling reaction times to probabilistically distributed events (Fig. 1A).

**Fig. 1.**
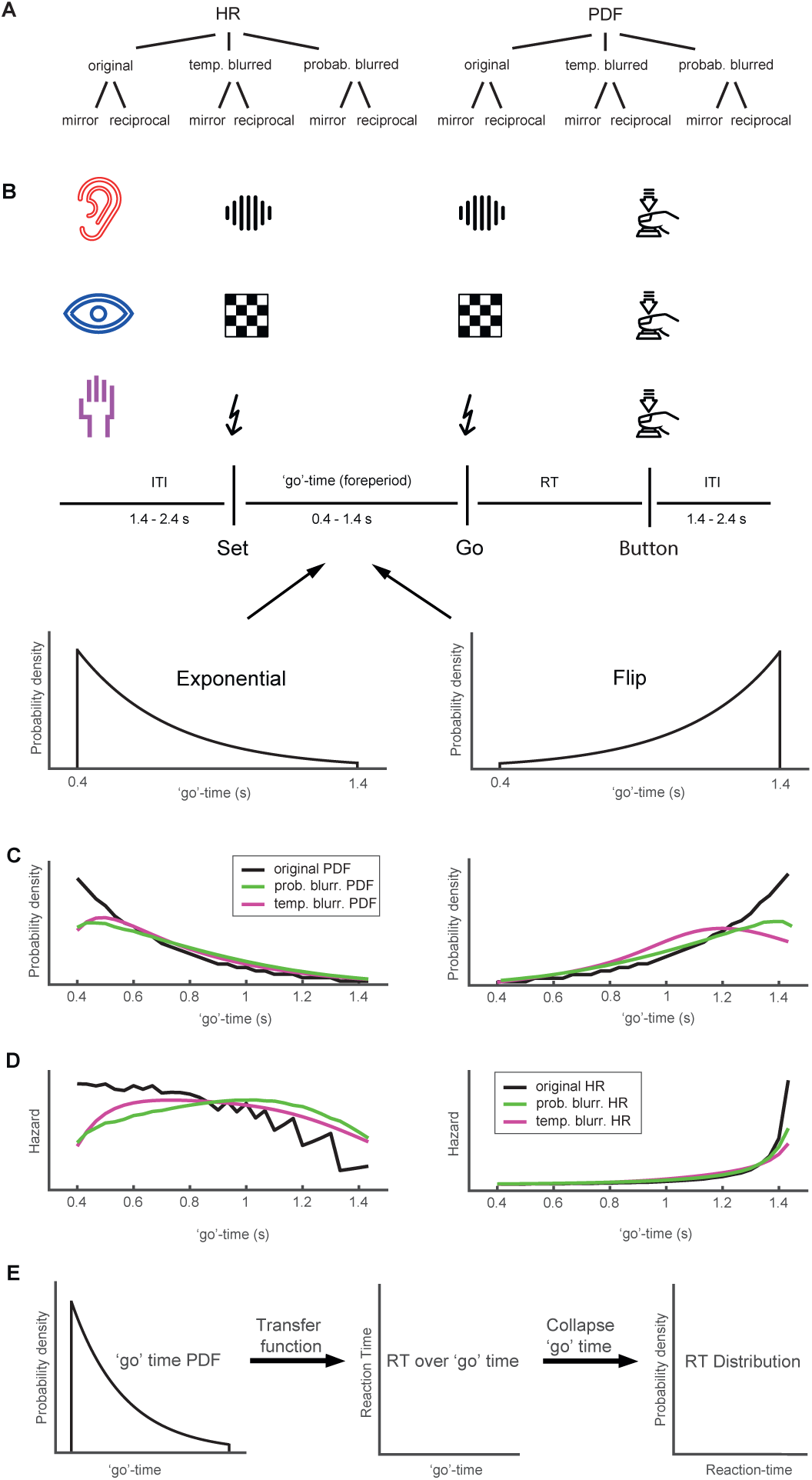
Models of event probability in time, task, and presented probability distributions. (A) Overview of variables for temporal anticipation. The canonical hazard rate (HR) model of temporal anticipation is based on three assumptions: 1) The brain employs the HR to anticipate events in time. 2) Time estimation contains uncertainty that scales linearly with elapsed time (*temporal blurring*). 3) Reaction time to events and the HR are linearly anti-correlated (*mirror*). The three assumptions are investigated in this paper using additional, PDF-based variables incorporating *probabilistic blurring* and a non-linear, *reciprocal*, relationship between RT and model (see Methods). (B) Schematic of ‘set’ – ‘go’ task. In auditory, visual, and somatosensory blocks of ‘set’ – ‘go’ trials, subjects were asked to respond as fast as possible to the ‘go’ cue. The time between ‘set’ and ‘go’ (the ‘go’ time) varied systematically. Within a block of trials the ‘go’ time was drawn from either a truncated exponential distribution (PD_Exp_) or its left-right flipped counterpart (PD_Flip_). (C) The two presented *event PDFs*, PD_Exp_ and PD_Flip_, in their original, temporally and probabilistically blurred versions (see Methods). (D) The hazard rates (HR) of PDF_Exp_ and PDF_Flip_ in their original, temporally, and probabilistically blurred versions (see Methods). (E, left arrow) The transfer function is a function of ‘go’ time and conceptualizes the mapping from event probability onto the corresponding reaction time. It reflects the brain’s represention of probability across time. (E, right arrow) The distribution of reaction times, irrespective of ‘go’ time, is used to identify both modality-specific and modality-independent processes.

These variables were derived from either of two basic parameters, namely the HR (*Hypothesis A*) or the PDF itself. In a first transformation, each basic parameter was subjected to one of two different types of blurring, reflecting uncertainty in the estimation of elapsed time. In the first type, *temporal blurring* (*Hypothesis B*), the variance of the Gaussian uncertainty kernel increases linearly with elapsed time. In the second type, termed *probabilistic blurring*, the variance of the kernel depends on the probability distribution of events across time (see Methods). The original, non-blurred, parameters were also tested in comparison to the blurred ones. Given the inverse relationship between reaction times and HR (*Hypothesis A*), in a second transformation all variables resulting from transformation one, were subjected to one of two types of inversion: either a linear transformation (“mirror”) which corresponds to mirroring a variable around its mean (*Hypothesis A*), or, for comparison, a simple nonlinear transformation, the *reciprocal* of a variable (see Methods). This reciprocal relationship between model and data implicates a nonlinear downweighting of low event probability (resulting in relatively longer RTs) relative to an upweighting of high probability (leading to relatively shorter RTs), which we hypothesize may reflect a more economic deployment of attention in time, benefitting the anticipation of event occurrence.

To investigate our hypotheses, we used a “set-go” paradigm (Fig. 1B), in which participants were asked to press a button with their right index finger as fast as possible in response to the ‘go’ cue. In 9.09 % of trials, no ‘go’ cue was presented (catch trial). In order to examine modality-specific and modality-independent aspects involved in event anticipation, the stimuli were presented separately in three sensory modalities (vision, audition, and somatosensation). The ‘go’ time (time between ‘set’ and ‘go’) was sampled from one of two different probability distributions, an exponential (PD_Exp_) and its “flipped” version (PD_Flip_). These two ‘go’ time PDFs (Fig. 1C, black lines) were selected because they are symmetric to each other, while their HRs are not (Fig. 1D, black lines). For each of these distributions, the different explanatory variables were derived and used to model the transfer function between each ‘go’ time and the corresponding RT. (Fig. 1E, left arrow). Differences and similarities between the three modalities were also studied at the level of RT distributions irrespective of ‘go’ time. (Fig. 1E, right arrow).

## Results

24 subjects performed experimental sessions on two consecutive days, generating approximately 3,500 RTs each. In all three modalities, the reaction times were strongly modulated by the ‘go’ time probability distributions. More specifically, the two symmetric PDFs presented, PD_Exp_ and PD_Flip_, lead to nearly symmetric patterns of RT. All explanatory variables were derived from the HR and PDF of each of the presented ‘go’ time distributions. First, we fitted the RT data with a linear model of the commonly proposed variable “temporally-blurred, mirrored HR”. Overall, this popular model failed to fit the data adequately. In the PD_Exp_ condition (Fig. 2A), the explanatory variable captured the behavior of the RT data mostly in the latter half of ‘go’ times. In the early part of the distribution, there were significant deviations between data and explanatory variable, reflected in the low R^2^ values of the fitted models.

**Fig. 2.**
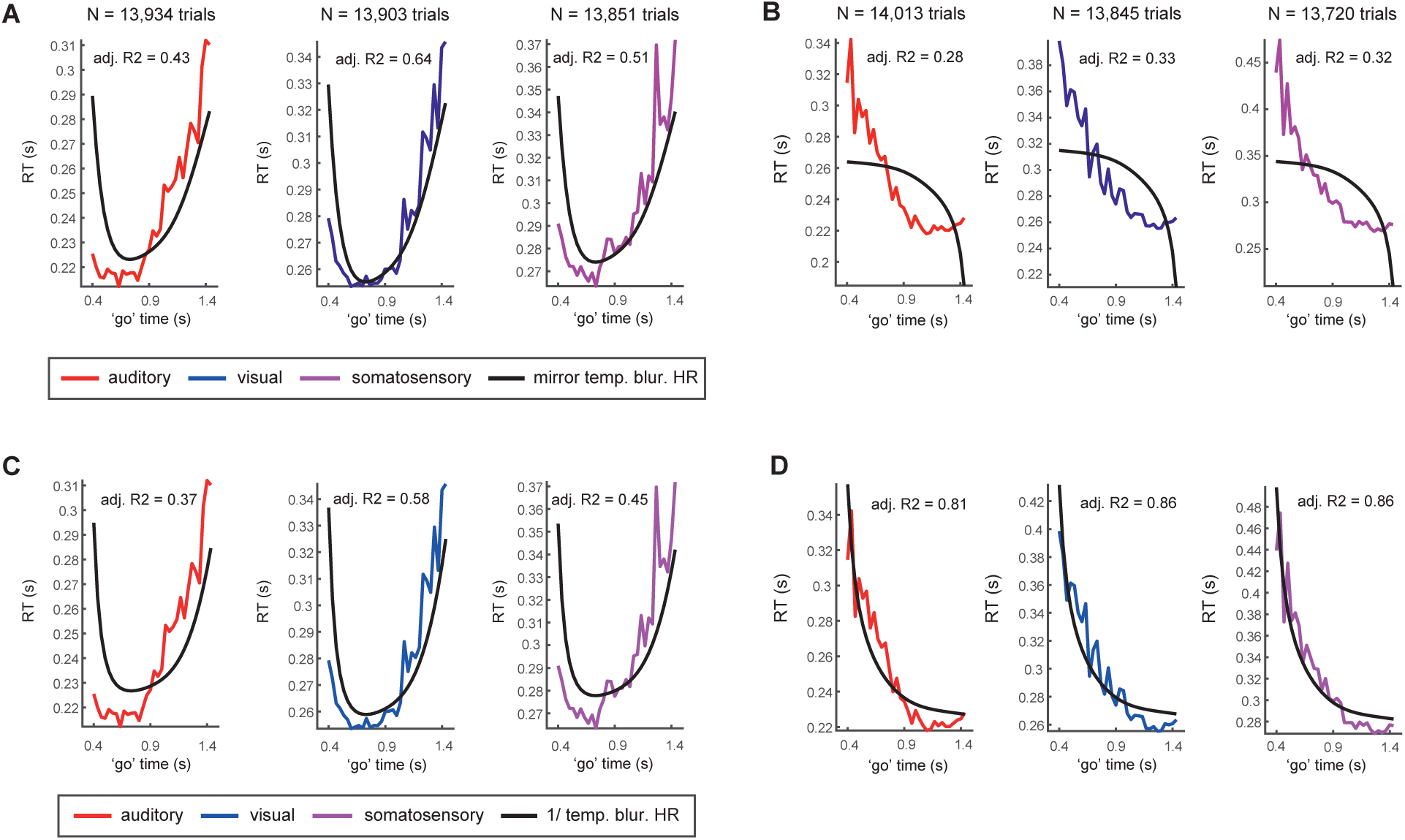
Models based on the temporally-blurred, mirrored hazard rate fail to capture reaction time data in exponential and flipped-exponential conditions. (A and B) The commonly used *temporally-blurred hazard rate* (black line) in its linearly transformed (“mirror”) version fitted to auditory (red), visual (blue) and somatosensory (violet) RT data from PD_Exp_ condition and from PD_Flip_ condition. (A) In the PD_Exp_ condition, the HR model deviates from RT in the early range of ‘go’ times. (B) Also note the substantial mismatch between mirrored HR and the RT in the PD_Flip_ condition. (C and D) Fits of the *temporally-blurred, reciprocal (“1/”) HR* to RT in the PD_Exp_ condition and in the PD_Flip_ condition. (D) Although the reciprocal relationship between model and RT improves the fit in the PD_Flip_ condition, the fit at the early ‘go’ times is not improved in the PD_Exp_ condition (C). See SI Appendix, Figs. S1 and S2 for a detailed summary of all other fitted hazard rate models.

In the PD_Flip_ condition (Fig. 2B), the deviation of the explanatory variable from the observed RTs was striking in all modalities, suggesting that *Hypotheses A* and *B* do not hold as canonical rules across different probability distributions. These results indicate that the mirrored HR is not employed by the brain in order to model event probability across time. We next tested whether the inverse relation between RT and HR can be better captured by a simple non-linear transformation. To do this, we replaced the mirrored transformation with the reciprocal, in which the HR is inverted by division instead of being linearly mirrored. This variable, the “temporally-blurred, reciprocal HR”, significantly improved the fit to RT in the PD_Flip_ condition in all modalities (Fig. 2D). However, in the PD_Exp_ condition we observed no improvement but rather a deterioration in the fit (Fig. 2C). Thus, the HR is an unlikely transformation for the brain to employ to model event probability across time.

### PDF-based models of RT capture data in three sensory modalities

We next turned to models based on a more fundamental, “core” probability parameter, the PDF itself. Similar to HR, we first examined the variable “temporally-blurred, mirrored PDF”. Although the shape of the explanatory variable captured trends in the data, including local minima which are near but not at the extrema of the ‘go’ time interval, the fit to RT was poor, especially in the PD_Exp_ condition (SI Appendix, Fig. S3C). We then examined the non-linear, reciprocal transformation by using the “temporally-blurred, reciprocal PDF” as an explanatory variable (Figs. 3A and B).

**Fig. 3.**
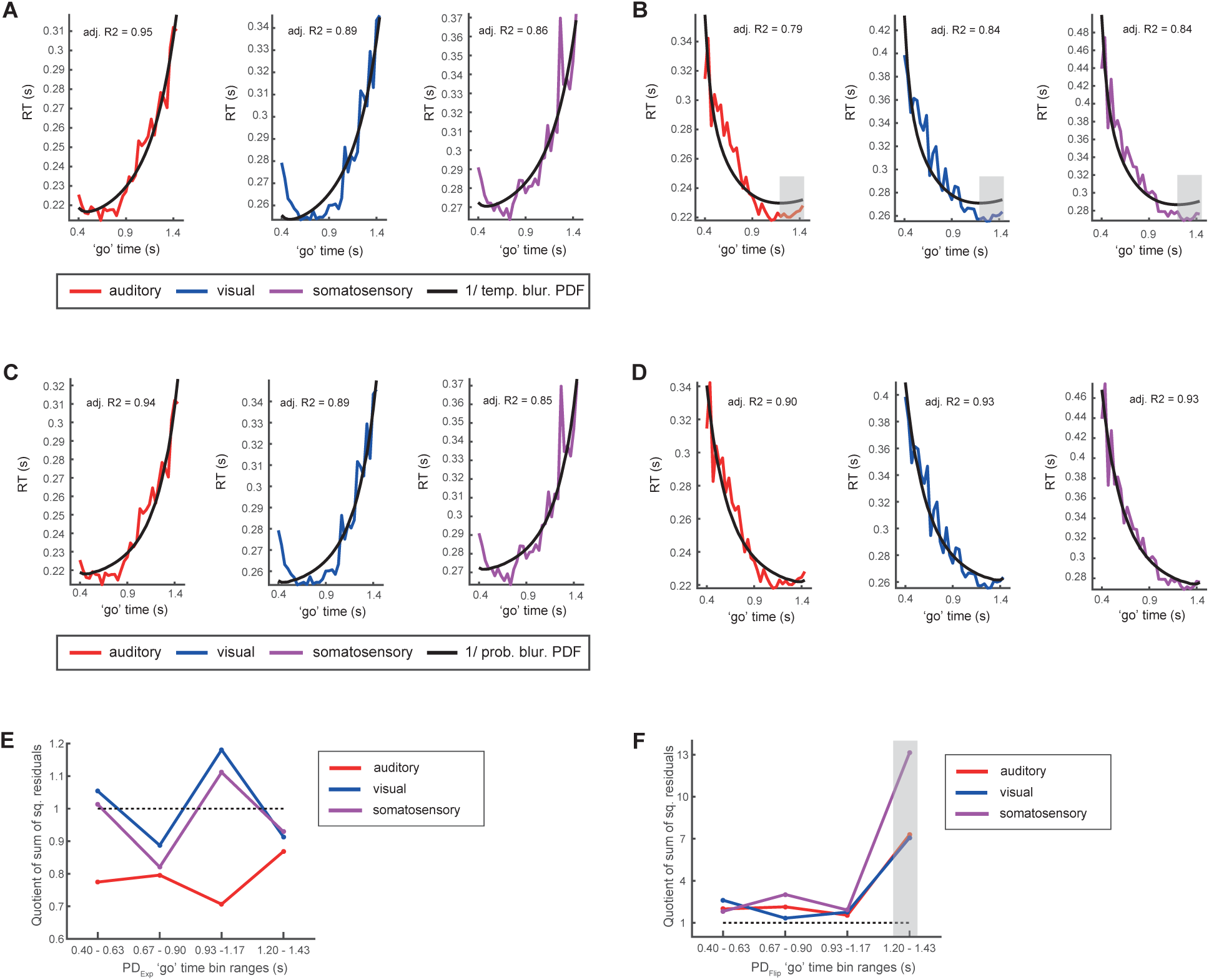
Models based on probabilistically-blurred, reciprocal PDF capture reaction time data in both exponential and flipped-exponential conditions. (A and B) Model fits to auditory (red), visual (blue) and somatosensory (violet) RT based on the *temporally-blurred, reciprocal PDF* (black line) in the PD_Exp_ condition (A) and in the PD_Flip_ condition (B). Since temporal blurring increases with ‘go’ time, it has a stronger smoothing impact on the model at longer ‘go’ times. This feature is not reflected in the data (‘go’ time span 1.2 to 1.43 s highlighted). (C and D) Model fits to RT based on the *probabilistically-blurred, reciprocal PDF* in the PD_Exp_ condition (C) and PD_Flip_ condition (D). (E and F) Comparison between two reciprocal PDF-based models (“temporally-blurred” and “probabilistically-blurred”), in each of four equally spaced bins across the ‘go’ time span. (E) In the PD_Exp_ condition, the quotient of squared residuals (“temporally-blurred” / “probabilistically-blurred”) is relatively close to 1 in all bins, indicating residuals of similar magnitude across the two models and thus a similar goodness-of-fit. (F) In the PD_Flip_ condition, residuals are bigger in the temporally-blurred model in all bins, especially in bin #4 (highlighted). See SI Appendix, Figs. S1 and S3 for a detailed summary of all other fitted PDF-based models.

The fit to the data was significantly improved, as shown by the similar behavior of model and data across ‘go’ time, which was confirmed by the high R^2^ values. Evidently, the reciprocal transformation of the PDF provided a much better fit than the mirror transformation as well as both transformations of the HR. In the PD_Flip_ condition in all modalities, we observed that the explanatory variable settles to a plateau of values quite early, at a ‘go’ time of around 1 s, while the actual data continue to have a negative slope (Fig. 3B, grey shading). Additionally, in the early range of ‘go’ times (e.g. 0.5-0.9 s) the explanatory variable decreases with a much steeper slope than the data. These systematic differences between model and data were introduced by the temporal blurring. Therefore, in the early range of ‘go’ times the Gaussian kernel has a small variance and causes only little blurring, less than the data suggest. In the late range of ‘go’ times, the Gaussian kernel has a large variance, leading to stronger blurring of the explanatory variable, much more than the data suggest.

### Uncertainty in elapsed time estimation is influenced by event probability and not only by elapsed time itself

These observations lead us to question the concept of monotonically increasing uncertainty that scales with elapsed time itself. Instead, we hypothesize that uncertainty in elapsed time estimation is modulated by temporal probability. This results in a blurring kernel with large variance in the early, less probable, range of ‘go’ times in the PD_Flip_ condition and in a blurring kernel with small variance in the late, more probable, range of ‘go’ times. We applied this *probabilistic blurring* to the reciprocal PDF and fitted this variable to RT. This simple operation drastically improved the fit in the PD_Flip_ condition (Fig. 3D). In the PD_Exp_ condition (Fig. 3C), the fit was almost identical to the temporal blurring case (Fig. 3A). This was to be expected, as in both blurring cases the variance of the Gaussian kernel monotonically increased with the magnitude of ‘go’ times. To quantify differences in model fit between temporally and probabilistically blurred variables, we divided the range of ‘go’ times into four equal-sized bins. Within each bin, the sum of squared residuals was calculated for the “temporally-” and “probabilistically-blurred, reciprocal PDF”. In the PD_Exp_ condition (Fig. 3E) the quotient is relatively close to 1 in all bins, indicating residuals of similar magnitude across the two models and thus a similar goodness-of-fit. In the PD_Flip_ condition (Fig. 3F) the quotient is generally larger, and always higher than 1, which shows that the probabilistically blurred model yielded smaller residuals in all bins and in all modalities. Clearly, probabilistic blurring represented the uncertainty in elapsed time estimation better than the commonly employed temporal blurring. For comparison and completeness, three more variables, the “probabilistically-blurred, mirrored HR”, the “probabilistically-blurred, reciprocal HR” and the “probabilistically-blurred, mirrored PDF” were also investigated (SI Appendix, Figs. S1-S3). None of these variables provided a better fit than the variable “probabilistically-blurred, reciprocal PDF”. We conclude that, in all three modalities the “*probabilistically-blurred, reciprocal PDF*” – but not the HR – is the most adequate model for mapping the brain’s temporal-probabilistic input onto its output, i.e. the reaction times.

### Probabilistically-blurred, reciprocal PDF captures RT to Gaussian distribution of events better than HR

The above results motivate the question whether our findings generalize to a different probability distribution. Therefore, we re-invited 18 subjects and presented them with a Gaussian distribution of ‘go’ times (see Methods). Again the “temporally-blurred, mirrored HR” could not adequately account for the data (SI Appendix, Fig. S4A). In contrast, the “probabilistically-blurred, reciprocal PDF” fitted the data in the auditory condition accurately (SI Appendix, Fig. S4B). The fits to visual and somatosensory data were less accurate, but in light of the simplicity of a model with only two free parameters, the PDF-based model captured the biphasic RT modulation.

### In a deterministic context, a probabilistically-blurred, reciprocal model of RT fits data better than HR

Up to this point, we investigated event anticipation in a *probabilistic* context, i.e. there is uncertainty about an event having occurred by the end of a timespan. This uncertainty is introduced by a percentage of catch trials which do not feature a ‘go’ cue. In contrast, in a *deterministic* context, an event will have occurred by the end of a timespan and accordingly, the uncertainty of event occurrence is zero. The HR is commonly suggested in both deterministic (5-8, 10) and probabilistic contexts (4, 9) as a model of RT. In order to investigate both the PDF-based and the HR-based variables’ potential to capture RT in a deterministic setting, we acquired data in auditory and visual sensory modalities using the same task and distributions (PD_Exp_ and PD_Flip_) but without catch trials. Clearly, the “temporally-blurred, mirrored HR” did not fit the data (SI Appendix, Fig. S5A and B), whereas the “probabilistically-blurred, reciprocal PDF” captured the data adequately in all conditions (SI Appendix, Fig. S5C and D). We conclude that in both probabilistic and deterministic contexts and in all investigated probabilistic conditions, the PDF-based model outperformed the HR-based model.

### RT is comprised of modality-specific and modality-independent components

The observed striking similarities in processing temporal-probabilistic structures between the three sensory modalities suggest shared neural processes, while the differences in, e.g., processing speed likely reflect modality-specificity. The RT distributions are the result of the superposition of such neural mechanisms. Accordingly, we characterize the RT distributions based on the hypothesis of separate, additive contributions to the mapping of an event PDF onto the corresponding RT (Fig. 1E). One contribution likely is *modality-specific*, reflecting more peripheral processing stages, e.g. latencies in sensory signal transduction and feed-forward information processing (21-28). The other contribution is hypothesized to reflect *modality-independent* processes associated more with the processing of the *‘go’ time PDF* itself. QQ-plots revealed overall similarities in *RT distribution* across PD_Exp_ and PD_Flip_ conditions, but also heavier right tails in the PD_Flip_ condition in all three modalities (Fig. 4A).

**Fig. 4.**
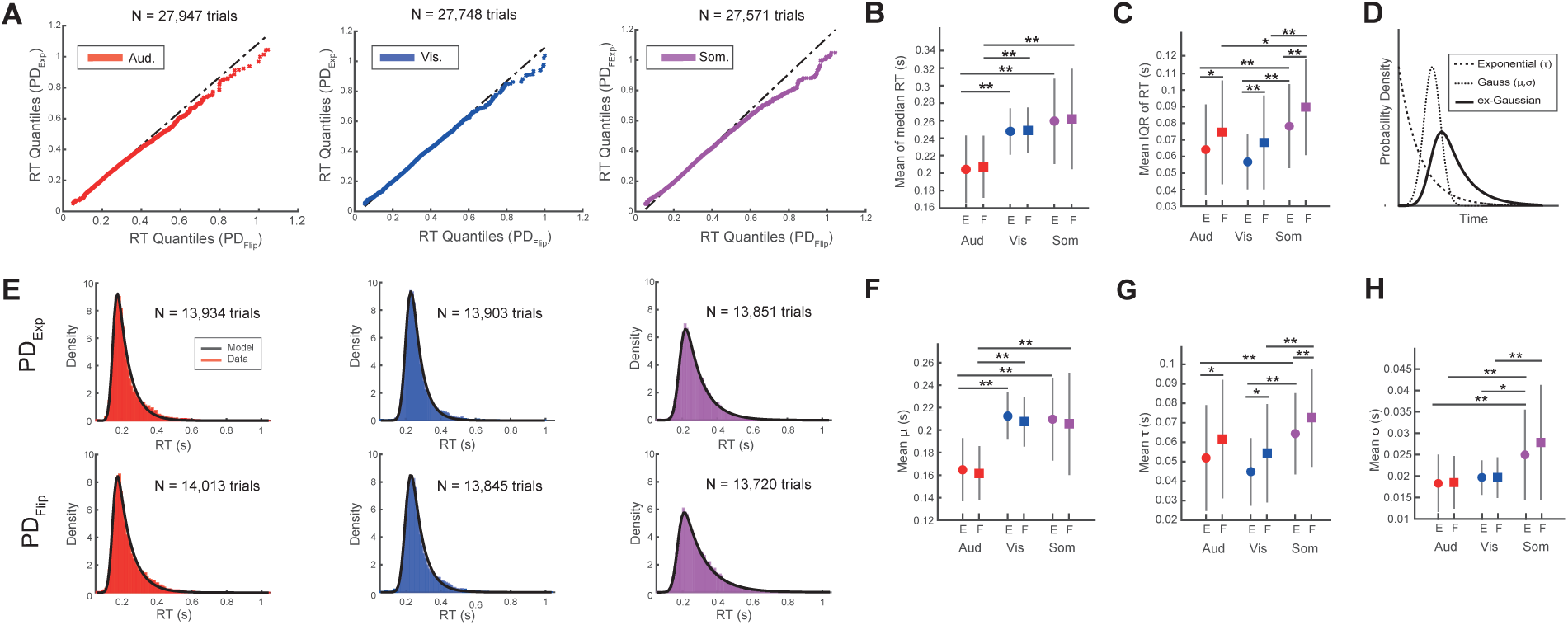
Reaction time variance is sensitive to the probability distribution of events. (A) In all three sensory modalities, QQ-Plots reveal heavier right tails in the reaction time distributions in the PD_Flip_ condition. (B) Mean of median RT in PD_Exp_ (‘E’) and PD_Flip_ (‘F’) conditions. Average RT is not sensitive to the presented probability distribution. (planned contrasts, * *P* < 0.05, ** *P* < 0.01, two-tailed Student’s *t*-test). (C) Mean of interquartile range of RT in PD_Exp_ and PD_Flip_ condition. RT variance is higher in the PD_Flip_ condition (planned contrasts, * *P* < 0.05, ** *P* < 0.01, two-tailed Student’s *t*-test). (D) The Ex-Gaussian distribution is a convolution of an exponential and a Gaussian PDF. (E) The ex-Gaussian model fits RT histograms on the group level (E) and on the single-subject level (see Figure S6). (F to H) Ex-Gaussian fit parameters from single-subject fits. (F) Mean of Gaussian *μ* resembles mean of median RT (compare to B, planned contrasts, * *P* < 0.05, ** *P* < 0.01, two-tailed Student’s *t*-test). (G) Mean of exponential *τ* resembles interquartile range of RT (compare to C, planned contrasts, * *P* < 0.05, ** *P* < 0.01, two-tailed Student’s *t*-test). (H) Mean of Gaussian *σ* is not sensitive to the presented probability distribution (planned contrasts, * *P* < 0.05, ** *P* < 0.01, two-tailed Student’s *t*-test). Error bars denote standard deviation. For ANOVAs, see SI Appendix, Tables S1 to S4.

We next studied these distributional differences. Participants’ RTs exhibited two characteristic patterns, one modality-specific, the other modality-independent. In the *modality-specific* pattern, median RT was faster in the auditory condition than in both visual (−42.4 ± 26.6 ms, *P* = 8.2*10^−7^; Tukey’s honest significant different test) and somatosensory conditions (−54.8 ± 33.7 ms, *P* = 1.1*10^−9^; Tukey) (Fig. 4B). In RT variance, a different *modality-specific* pattern emerged. Interquartile range (IQR_RT_) was smaller in vision (−21.3 ± 26.4 ms, *P* = 0.00026; Tukey) and audition (−14.6 ± 26.2 ms, *P* = 0.0198; Tukey) compared to somatosensation (Fig. 4C). In the *modality-independent* pattern, the change in *‘go’ time PDF* had no effect on median RT (Fig. 4B). However, IQR_RT_ was significantly larger in the PD_Flip_ condition as compared to PD_Exp_ (Fig. 4C). The magnitude of the difference in IQR_RT_ between probabilistic conditions did not differ between sensory modalities (*F*_(2,22)_ = 0.04, *P* = 0.96, one-way ANOVA (for analysis of variance and planned contrasts on median RT and IQR_RT_ see Tables S1 to S4.). To identify how these systematic differences relate to the observed *modality-specific* and *modality-independent* components of the RT distributions, we modeled the RT distributions with an exponential-Gaussian PDF. This two-process model is the convolution of an exponential PDF (parameter *τ*) and a Gaussian PDF (parameters *μ* and *σ*) (Fig. 4D). It states that the generation of RT depends on a sum of peripheral Gaussian processes and a central decision process that is hypothesized to be exponential (14, 29). The model provides an excellent fit to the data in all conditions, both at the group level (Fig. 4E) and at the single-subject level (SI Appendix, Fig. S6). Both Gaussian parameters, *μ* and *σ*, were *sensitive* to changes in sensory modality but were *insensitive* to changes in the *‘go’ time PDF* (Figs. 4F and H), which is in line with the model’s implicit claim of peripheral Gaussian processes. In contrast, the exponential parameter *τ* was *sensitive* to the *‘go’ time PDF* (Fig. 4G), having larger values in the PD_flip_ condition as compared to PD_Exp_. This pattern of *τ* closely mirrors the behaviour of IQR_RT_ (Fig. 4C) (See SI Appendix, Tables S1-S4 for analysis of variance and planned contrasts on *μ, σ*, and *τ*). We found that Gaussian *μ* captured the behavior of median RT in all three modalities (SI Appendix, Fig. S7A and B). Likewise, and also independent of modality, *τ* captured IQR_RT_ (SI Appendix, Fig. S7C and D). Taken together, these findings suggest that only the process(es) reflected in the exponential parameter *τ* use information on the *‘go’ time PDF*. Taken together, our findings on the influence of the *‘go’ time PDF* on the *RT distribution* support (i) the broader hypothesis of peripheral and central processes involved in event anticipation as well as (ii) the specific claim of a central exponential process shared by all three modalities.

## Discussion

We investigated the brain’s capabilities to infer probability as a function of time based on sensory input, by analysing reaction times to temporally distributed events. Subjects performed auditory, visual, and somatosensory ‘set’ – ‘go’ tasks in which the time between the two events was drawn from different PDFs. In all modalities, subjects clearly extracted and used the probabilistic information encoded in the trial structure to predict ‘go’ cue onset, as shown by their distinct patterns of RT modulation. Previous work has proposed the hazard rate as a model of probability over time used by the brain to anticipate events and plan responses. We demonstrate that it is not the HR but the PDF – a simpler and more stable variable – which better captures the data. We argue that these findings represent a significant contribution towards a mechanistic understanding of long-standing questions regarding how the brain extracts its environment’s temporal structure to adapt behavior.

We found that RT was sensitive to changes in event probability density. Notably, the observed RT modulation in the PD_Exp_ condition indicates that subjects inferred temporal-probabilistic information from exponentially distributed events. This is remarkable because it is commonly assumed that an exponential distribution renders temporal prediction impossible (7, 30), arguably due to its flat HR. Interestingly, the two symmetric ‘go’ time PDFs lead to near-symmetric patterns of RT. The similarity in RT temporal modulation across the three sensory modalities suggests that similar aspects of the underlying representation of the temporal-probabilistic structure are used by the brain to model and predict stimulus onset in each modality. The specific nature of this model has been a subject of debate in the literature, with the hazard rate being argued to play a central role (3-6, 8-10, 13). In particular, it has been proposed that the brain’s model of event probability across time is linearly anticorrelated with the HR (*Hypothesis A*). An additional prominent assumption of this model is that uncertainty in elapsed time estimation increases monotonically with time itself (*Hypothesis B*).

We first modeled the RTs using various explanatory variables based on linear and nonlinear transformations of the HR. None of these variables adequately captured the behavior of RTs across the different ‘go’ time distributions. Therefore, *Hypothesis A* does not hold as a canonical rule, and the hazard rate is not a likely transfer function the brain uses to relate an event PDF to its corresponding RT. Instead, we demonstrate that a nonlinear transformation of the PDF, the reciprocal, better captures the RT behavior across the different PDFs in all three sensory modalities.

Interestingly, the nonlinear, reciprocal transformation of the temporally-blurred HR improved drastically the fit to RT as compared to the linear transformation of HR (compare Figs. 2B and D). This shows that a nonlinear relationship between event probability and corresponding RTs yielded a better model fit than a linear relationship, even in the case of the HR. Examining the equation for the reciprocal HR, it is easy to see that the reciprocal HR is just the reciprocal PDF multiplied by the survival function, as 1 / HR = (1 – CDF) * 1 / PDF. In this equation the survival function appears as a time-varying scaling factor of the reciprocal PDF.

The models based on the reciprocal PDF clearly provided the better fits to RT in all conditions as compared to the reciprocal HR. Following this revealing result that the PDF, and not the HR, is the most likely parameter modeled by the brain, and given that HR = (1/survival function) * PDF, it is obvious that the brain did not employ the scaling factor (1/survival function) in order to scale the PDF. Instead the scaling appeared to be better approximated by a fixed value, uniform for the entire range of go-times. Nonetheless, it seems possible that in contexts other than simple event anticipation, the brain might use the survival function as a scaling factor for event probability density in which case the HR may be an appropriate parameter of probability in time.

We note that in a *reward*-based context, the HR has been shown to adequately describe temporal expectation (4-7, 10) in a 100 % rewarded condition, but not when uncertainty of reward is introduced (30, 31). In the experiments we performed here, no reward was delivered. Therefore, the fact that the PDF, but not the HR, provided the best model of RT might be related to the absence of reward. Although it is clearly beyond the scope of our study to identify how reward modulates anticipation, a simple, intuitive hypothesis can be formulated. If the brain employed a scaling factor for the PDF, which in our case is uniform across time, then under reward conditions this scaling factor could approximate 1/(survival function), and the resulting parameter encoded would be the HR, as HR=(1/survival function)*PDF. The survival function is defined as 1-CDF, where the CDF captures the cumulative probability that an event should have happened up to and including the current time instance. The computation of this accumulated probability is cognitively demanding and would likely be facilitated by motivating effects of expected reward. This simple hypothesis could describe a basic mechanism by which the brain incorporates reward into event anticipation (SI Appendix, Fig. 8A). It suggests a possibility how the PDF-based model we describe here could link to the HR-based models in the reward literature.

Commonly, HR-based models of reaction time incorporate the concept of *temporal blurring* (3, 5, 6) which reflects properties of scalar timing, i.e. the brain’s uncertainty in elapsed time estimation increases linearly with time (*Hypothesis B*) (18): longer intervals carry higher uncertainty in their estimation than shorter intervals. This implies that the brain’s capacity to react fast and accurately to longer timespans is limited compared to shorter timespans, irrespective of the accuracy of the brain’s estimate of event probability in time. In other words, the error in time estimation ultimately constrains the brain’s benefit from temporal-probabilistic inference. By modeling RT, we found that probabilistically-blurred models were superior to the temporally-blurred ones. In particular, the RTs at the most probable, longer ‘go’ times in the PD_Flip_ condition were much better captured by the probabilistically-blurred model compared to the temporally-blurred one (Figs. 3B and D). This finding challenges the common assumption that the brain models elapsed time with uncertainty that increases with time *per se (2)*. Instead it seems that uncertainty in time estimation also scales with probability. We suggest that by modeling its environment’s temporal-probabilistic structure, the brain can overcome what has been considered an built-in limitation: the uncertainty in time estimation.

In addition to event probability in time and uncertainty in time estimation, a third source of uncertainty can be quantified in event anticipation: the uncertainty of event occurrence. Suppose a ‘go’ cue will certainly have occurred by the end of a trial. In this case there is no uncertainty of event occurrence towards the right extremum of the ‘go’ timespan. This is defined here as a *deterministic* context. In contrast, ‘go’ cue occurrence may remain uncertain when in a percentage of trials no event occurs (catch trials), which we define as a *probabilistic* context. The HR has been proposed in both *deterministic* (5-8, 10) and *probabilistic* contexts (4, 9) as an important model of probability in time. Notably, in a *deterministic* setting, certainty of event occurrence is reflected by the CDF asymptotically approaching 1. This leads to a steep increase of the HR’s slope towards the end of the ‘go’ time period, irrespective of probability density (SI Appendix, Fig. S8B) – the exponential PDF being a rare exception to this. Commonly, this CDF-based up-weighting of ‘go’ time probability is argued to reflect anticipation, which conforms with intuition as it maximizes event probability towards the right extremum of a timespan when the event will inevitably occur. However, as a result of the CDF approaching one, the HR’s slope becomes very steep and its values approach infinity. Since there is a lower bound to reaction time, this behavior towards the end of a timespan challenges the concept of the HR as a model of RT. Although the commonly employed temporal blurring remedies this problem somewhat by reducing HR values as time increases, the conceptual issue of an ever-increasing variable remains. Another general problem of the HR as a canonical model of RT in event anticipation concerns its instability in calculation in both *deterministic* and *probabilistic* contexts. The HR is difficult to estimate from empirical data because technically, it requires several steps: computation of PDF, integration of PDF to arrive at CDF, transformation of CDF to arrive at the survival function, division of PDF by survival function. Even small errors in the representation of PDF or its CDF will lead to large and unpredictable errors in HR which could have considerable consequences for an organism relying on the HR as its model of temporal probability. The term 1 / PDF, on the other hand, can be interpreted as “one-in-many” – a much simpler and more stable computation. For example, if the probability of an event is 0.02, the brain might interpret it as “1/0.02”, i.e. “one-in-fifty”. We also note that in some of the previous work on the topic, the observed relation between RT and HR might also have been well-captured by the PDF, as both HR and PDF can be monotonically in- or decreasing.

Taken together, we demonstrated that in a *deterministic* setting, the probabilistically-blurred, reciprocal PDF captured the RT data better than the HR-based models. Although the addition of a small percentage of catch trials, which created a *probabilistic* context, reduced the HR’s extreme behavior at the right extremum of the PDFs’ timespan, we again found that the PDF-based model fitted the data better than the commonly proposed HR-based models.

To investigate the influence of sensory input modality on processes involved in temporal-probabilistic inference, we analysed the full RT distributions. We observed modality-specific differences in average RT that are in agreement with existing findings covering a wide range of simple RT and go/no-go tasks (14, 32). Here they are seen in the context of event anticipation. Also in agreement with previous work, we found modality-specificity in RT variance, which is a measure of the temporal precision of the response (14). Interestingly, in all three sensory conditions, variance but not average RT was sensitive to event probability density.

The RTs could be decomposed into the sum of a *modality-independent* PDF which was *exponential* and a second, *modality-dependent* PDF which was *Gaussian*. The Gaussian parameter *μ* corresponded to the modality-specific offsets in average RT. The Gaussian parameter *σ* displayed a different modality-specific pattern. Neither Gaussian parameters *μ* nor *σ* were sensitive to changes in the event PDF of the input. We conjecture that the Gaussian part of the ex-Gaussian model represents *modality-specific* processes of a peripheral nature, e.g. the contingencies of the sensory apparatus, i.e. signal transduction and conduction, and also shared motor processes. In contrast, the *exponential* part of the ex-Gaussian model was sensitive to the event PDF. It also captured the difference in RT variance between the two probability distributions in all three sensory modalities. This outcome is in agreement with proposed theories in which the exponential part corresponds to a central process. (14, 29). Our findings suggest all three sensory modalities share a similar exponential process – which in turns makes specific predictions for what neural activity patterns should be sought on recordings from the relevant sensory and supra-sensory regions.

In summary, there are both modality-specific and modality-independent processes involved in the modeling of temporal probability by the brain. RT in a simple RT task can be decomposed into two components, one controlled by modality-dependent factors and the other independent of modality. The latter component, however, changes to reflect the modality-independent objective distribution of ‘go’ times selected by the experimenters. The modality-specific processes can be confidently hypothesized to reside within the neural substrate mediating each individual modality. However, from these behavioral data it is not possible to conclude whether the modality-independent processes have a central implementation, shared by all modalities, or whether they reflect the same canonical computation within the neural substrate of each separate sensory modality. We expect that these findings will aid efforts in the understanding of the neural mechanisms involved in predictive processes. Our results demonstrate that irrespective of sensory input modality, the brain models its environment’s temporal-probabilistic structure using a nonlinear transformation of the PDF, but not the hazard rate.

## Material and Methods

### Ethics statement

The experiments were approved by the Ethics Council of the Max-Planck Society. Written informed consent was given by all participants before the experiment.

### Subjects

24 human subjects (13 female), aged 19-33, participated in the auditory, visual, and somatosensory experiments. An additional 18 subjects (13 female), aged 19-33 participated in an auditory and visual control experiment (“no-catch-trials experiment”). All were right-handed and had normal or corrected-to-normal vision and reported no hearing impairment and no history of neurological disorder. Participants were naive to the purpose of the experiment. They received €10 per hour for participating.

### Task and procedure

In auditory, visual, and somatosensory conditions, subjects performed a simple ‘set’ – ‘go’ task in which a ‘set’ cue was followed by a ‘go’ cue. The timespan between the onset of both cues, termed the ‘go’ time, was a random variable that was drawn from a specific probability distribution. Subjects were asked to foveate a central black fixation dot and respond as fast as possible to the onset of the ‘go’ cue with a button press on a response device using the right index finger. After a button press, a small black circle appeared for 0.2 s around the central fixation dot indicating the end of the trial. In some trials, no ‘go’ cue appeared (‘catch trials’), in which case participants were instructed to not press the button. In these catch trials, a small black circle appeared 1.9 s after ‘set’ cue onset, indicating again the end of the trial. The experiment consisted of two separate sessions taking place at the same time of the day on two consecutive days. A single session consisted of four blocks per sensory modality (vision, hearing, touch) and lasted approximately 2.5 hours. Per block, 165 trials were presented (including 15 ‘catch trials’), resulting in 1,980 trials per session for each subject (3,980 trials for two sessions), and a total of 95,040 trials for all subjects. A short training block was run before the first block of each sensory modality on both days to familiarize subjects with the task. During all experimental blocks, subjects wore headphones and positioned their heads on a forehead-and-chin rest (Head Support Tower, SR Research Ltd.) at a fixed distance of 60 cm relative to the computer monitor. Each subject’s dominant eye, as determined by Miles test (33) was tracked at a sampling frequency of 1,000 Hz (Eyelink DM-890, SR Research Ltd.). Subjects were asked to restrict eye blinking to the timespan after a button press, i.e. during the ITI. Trials in which visual fixation was not maintained within a radius of 2.5 degrees visual angle around the central fixation point for more than 300 ms during the ‘go’ time were automatically discarded for data analysis. All stimuli were generated using MatLab (The MathWorks, Natick MA, USA) and the Psychophysics Toolbox (34) on a Fujitsu Celsius M730 computer running Windows 7 (64 bit). The experiment took place in a dimly lit, soundproof booth.

### Auditory stimuli

Two white noise bursts (50 ms duration, 8 ms cosine ramp, onset and offset) served as ‘set’ and ‘go’ cues. They were presented at 60 dB SPL above hearing threshold at 1 kHz, as determined by pure tone audiometry according to ISO 8253. All auditory stimuli were output by a high-quality interface (RME Fireface UCX) and delivered diotically using electrodynamic headphones (Beyerdynamic DT 770 PRO) driven by a headphone amp (Lake People GT-109). The sound pressure level was calibrated to 75 dB(A) individually for each transducer while using a temporal weighting suited for impulsive stimuli (/*τ* = 35ms). To this end we used an IEC 603184 artificial ear simulator (model G.R.A.S. 43AG) with according pinnae and a IEC 60942 class 1 sound level calibrator (Larson Davis CAL200) and an IEC 942 class 1 pistonphone with barometric correction as calibration source (G.R.A.S. Type 42AA).

### Visual stimuli

The visual ‘set’ cue (duration 50 ms) consisted of two checkerboard patterns which were presented simultaneously. One was positioned in an upper quadrant, the other in the lower quadrant on the opposite side. The ‘go’ cue consisted of two checkerboard patterns the same location but the with black-white pattern reversed. Each checkerboard subtended 6.6 × 6.6 degrees of visual angle and consisted of 7 × 7 black and white squares, each subtending 0.9 × 0.9 degrees of visual angle. The center of each checkerboard was positioned at a horizontal distance of 12.6 degrees of visual angle and at a vertical distance of 7.1 degrees from the center of a central, black fixation dot. The site of presentation alternated between the left and right side on a trial-by-trial basis. Visual stimuli were presented on a BenQ XL2420-B monitor (resolution 1,920 × 1,080, refresh rate 60 Hz) which was set to a gray background.

### Somatosensory stimuli

Two short electric pulses (duration 200 µs) were presented as ‘set’ and ‘go’ cues using a constant current stimulator (Digitimer DS7A). Each subject’s perceptual threshold was determined by increasing stimulus intensity (mean current) until the subject first reported a sensation and then decreasing it until it was no longer perceived. The perceptual threshold was recorded as the lowest ascending stimulus intensity at which the subject reported sensation. For the experimental task, the electric current was set to a higher intensity (mean current = 7.9 +/- 3.9 mA) that the subject judged comfortable, yet easily perceptible.

### Temporal probabilities

The ‘go’ time was a random variable drawn from one of two probability distributions (Fig. 1B) that was fixed throughout two consecutive blocks of trials. The distributions were chosen by parametrically searching the family of Weibull distributions for cases that would fulfill the three following criteria:

1. The one distribution should be the left-right flipped version of the other so that this symmetry in PDFs would help to identify the effect of elapsed time itself on the modulation of RTs by the probability distributions.
2. The two distributions should have HRs with opposite slopes, in order to investigate the modulation of RTs by HR.
3. The number of trials in each of the quintiles of the probability distribution should be similar. This criterion was selected so that the modulation of variance of RTs across the ‘go’-time interval can be investigated while keeping the number of trials used for the estimation of variance roughly the same. As the probability distributions only contained integer values (i.e. the number of trials for each ‘go’-time) it was not possible to identify PDFs with exactly the same number of trials in each quantile. This criterion was relaxed so that each quintile should have the same number of trials ± 2.5 % with the neighboring quantiles.

A parametric search identified a Weibull distribution with parameters *k* = 1 and *l* = 0.33:

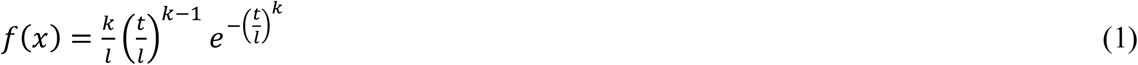

The identified shape value *k* =1 reduces the Weibull to an exponential distribution:

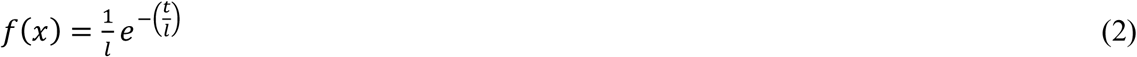

The x-axis of both distributions was discretized with the step size of the monitor refresh rate. This is based on the assumption that this level of discretization is not perceivable in the context of the ‘set’ – ‘go’ design, rendering the distributions continuous to the brain. The y-axis was also discretized as the PDF described number of trials at each discrete ‘go’ time point. Both distributions were delayed by 0.4 s giving a range of ‘go’ times from 0.4 to 1.4 s. To minimize sequential effects, the ‘go’ times were randomized with the constraint that no more than two consecutive trials had the same ‘go’ time. The intertrial interval (ITI, range 1.4 to 2.4 s) was randomly drawn from a uniform distribution. During each session in auditory, visual, and somatosensory conditions half of the blocks followed the exponential distribution (PD_Exp_), the other half followed its flipped counterpart (PD_Flip_). The probabilistic structure changed after two blocks without notification. To control for order effects, the conditions (sensory modalities and probability distributions) were organized in a Latin square design, based on which modality and distribution were shuffled across subjects and days.

### Gaussian ‘go’ time distribution

To investigate event anticipation in a probabilistic setting different from the exponential and flipped exponential ‘go’ time distributions, 18 subjects were reinvited and participated in a third experimental session. The subjects performed auditory, visual, and somatosensory ‘set’ – ‘go’ trials as described above, which included 9.09 % catch trials. Following the procedure outlined above, the time between ‘set’ and ‘go’ was drawn from a Gaussian distribution with parameters *μ* = 0.9 and *σ* = 0.25. The distribution was truncated at the flanks, giving a range of ‘go’ times from 0.4 to 1.4 s, giving the distribution a spread of two standard deviations around the mean. To minimize sequential effects, the ‘go’ times were randomized with the constraint that no more than two consecutive trials had the same ‘go’ time. The intertrial interval (ITI, range 1.4 to 2.4 s) was randomly drawn from a uniform distribution. To control for order effects, the order of sensory conditions was shuffled across subjects. Each subject generated 300 reaction times per sensory modality.

### Control experiment without catch trials

To investigate event anticipation in a *deterministic* setting, where every trial contains a ‘go’ cue, a control experiment was conducted. Using the same ‘set’- ‘go’ task as described above, auditory and visual blocks of trials were presented. Half of the blocks followed the exponential distribution (PD_Exp_), the other half followed its flipped counterpart (PD_Flip_). No catch trials were presented. To minimize sequential effects, the ‘go’ times were randomized with the constraint that no more than two consecutive trials had the same ‘go’ time. The intertrial interval (ITI, range 1.4 to 2.4 s) was randomly drawn from a uniform distribution. The probability distribution changed after two blocks without notification. To control for order effects, the conditions (sensory modalities and probability distributions) were organized in a Latin square design, based on which modality and distribution were shuffled across subjects. A new group of 18 subjects was invited and performed the task. Per sensory modality, each subject produced 240 reaction times under each probabilistic condition, which sum up to a total of 920 reaction times per subject for two modalities and two probabilistic conditions.

### Exponential-Gaussian model

To quantitatively investigate the distributional properties of RT between conditions, we used the exponential-Gaussian distribution as a well-established parametric two-process model of RT (14, 29, 35, 36).

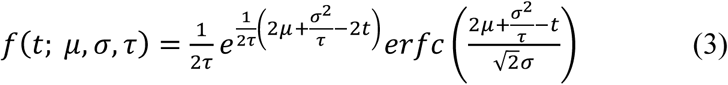

where *erfc* is the complementary error function defined as

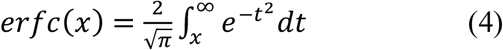

The parameters *μ* and *σ* are the mean and standard deviation respectively of the Gaussian constituent of the ex-Gaussian distribution. Parameter *τ* is the exponent of the exponential constituent distribution.

The best-fitting Ex-Gaussian parameters were obtained for each subject in each condition using a least-squares fitting algorithm. We observed no systematic difference in adj. R2 between conditions (SI Appendix, Fig. S6B). Therefore the ex-Gaussian proved to be a well-fitting model of RT in all conditions.

### Temporally blurred (‘subjective’) PDFs

The PDFs were blurred by a temporal uncertainty kernel that scales with elapsed time from a reference time point. More intuitively, the longer the elapsed interval to be estimated, the bigger the uncertainty about its length. In recent work it has been hypothesized that this uncertainty kernel has a Gaussian shape and its standard deviation increases linearly with time as *σ*= *φ* · *t*, where *t* is the elapsed time and *φ* is the scale factor by which the standard deviation *σ*of the Gaussian uncertainty kernel increases (5). This intuitively implies that the ratio of the standard deviation of temporal uncertainty to the elapsed time is constant and for this reason the variable *φ* has been termed a Weber fraction of the estimation of elapsed time (5).

Each of the employed distributions is characterized by the following three functions

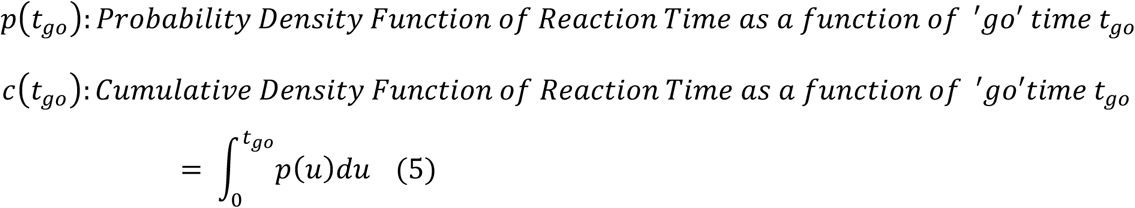

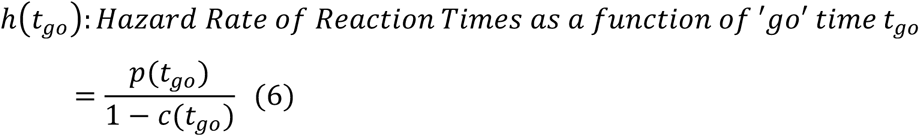

Each of the PDFs were blurred with a Gaussian kernel with variance increasing with time. The equations for the corresponding subjective functions are:

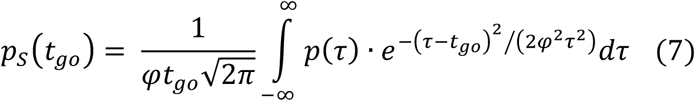

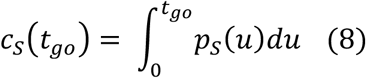

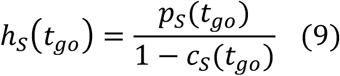

From equation (7) it is evident that for a given ‘go’ time *t*_*go*_ the PDF is convolved with a Gaussian kernel centered at *t*_*go*_.

The distributions used in this work were not continuous but discrete. They were only represented at the possible time points of stimulus presentation in either auditory, visual or somatosensory conditions. The stimuli presentation instances for PD_Exp_ and PD_Flip_ ranged between [0.4 1.4] sec at steps of 2/60 s (every 2 frames with frame rate 60 frames per second). The computation of the CDF from the discrete PDF was performed using trapezoidal integration. The hazard rate was computed from these discrete versions of a PDF and a CDF. The relatively small sampling interval resulted in discrete CDFs and hazard rates closely approximating the expected continuous versions of these functions from the analytical solutions (SI Appendix, Fig. S9).

The shortest ‘go’ time is 0.4 sec after ‘set’ cue onset. At this time point the uncertainty kernel has standard deviation *φ* · 0.4. Similarly at the longest ‘go’ time of 1.4 sec this uncertainty has standard deviation *φ ·* 1.4. In order to implement equation (7) for the computation of the subjective PDF, the definition of the PDF was extended to the left and right of the actual stimulus presentation interval as:

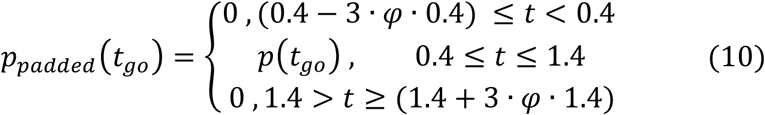

The extensions were equal to 3 standard deviations of the Gaussian uncertainty function at the shortest and longest ‘go’ times. This 3-standard-deviation extension was selected as it encapsulates the 99.7% of the Gaussian uncertainty. Then the integral in equation 3 was computed between these new extrema [(0.4 – 3 · *φ* · 0.4), (1.4 + 3 · *φ* · 1.4)] instead of the impractical interval of minus to plus infinity.

The selection of the value of *φ* was based on previous research (37, 38). With this value of *φ* = 0.21 the temporal range of the extended PDF as defined in equation (10) becomes [0.148 2.28] s which is also the range of integration in the computation of the subjective PDF in equation (7).

The PDF of each distribution, *p*(*t*_*go*_) was normalized so that its integral does not amount to one but to 0.9091, which is the total probability that an event will occur over the timespan covered by the distribution. The remaining 9.09 % of trials (30 out of 330) are catch trials in which no ‘go’ cue was presented. Consequently the maximum value of the resulting CDF *c*(*t*_*go*_) is 0.9091. The catch trials introduced some uncertainty about whether the event will occur at all. The function of the catch trials in the context of modeling temporal-probabilistic structures is to prevent the brain’s estimation of a PDF from being confounded by the expectation of a conditional event, i.e. the mandatory occurrence of a ‘go’ cue at the end of a given ‘go’ time range. It is easy to see from equation (6) that the introduction of catch trials and the corresponding reduction of maximum CDF to a value significantly smaller than one stabilizes the computation of the hazard rate at the right extremum of the PDF, i.e. the part where the CDF would approach one in the absence of catch trials. Here it has to be mentioned that although the maximum CDF value is 0.9091, the denominator in the computation of the HR remain 1 – *c*(*t*_*go*_) and not 0.9091 – *c*(*t*_*go*_). This is because this denominator represents the probability that nothing has happened up to time point *t*_*go*_. This term includes also the probability that nothing has happened up to *t*_*go*_ because the current trial is a catch trial. So this denominator could be alternatively defined as *c*_*catch*_ + *c*_*max*_ – *c*(*t*_*go*_), where *c*_*max*_ is the maximum CDF value of the stimulus PDF, equal to 0.9091, and *c*_*catch*_ is the probability that no ‘go’ cue appears at all, which is equal to 0.909. As these two terms sum to one, the denominator in the computation of the HR is correctly stated as in equation (6).

### Probabilistically blurred PDFs

In the definition of the subjective function in equation (7), the PDF was convolved with a Gaussian function, which represented the uncertainty in elapsed time estimation at a given time point. This uncertainty has been hypothesized to increase linearly with time (1), irrespective of the probability density function of event occurrence. In addition to this, here an alternative hypothesis was investigated in which the uncertainty in elapsed time estimation depends on the probability density function of event occurrence. The hypothesis states that ‘go’ times with high probability of event occurrence are associated with low uncertainty in time estimation based on the rationale that the brain predicts the onset of upcoming events of high probability more accurately. In contrast, ‘go’ times with low probability carry higher uncertainty, even if the time they span is short.

This probabilistic blurring of elapsed time estimation was implemented in a similar fashion to the temporal blurring described in equation (7). However the standard deviation of the Gaussian kernel does not scale linearly with time as in the temporal blurring case (*σ* = *φ* · *t*), instead it scales according to the PDF of event occurrence. In order to use realistic variance values in the blurring Gaussian kernel the minimum and maximum values of the standard deviation were set accordingly to the temporal blurring case as

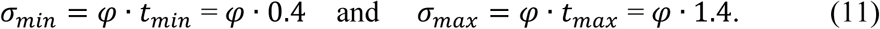

The PDF under investigation was then scaled so that its minimum value is *σ*_*min*_ and its maximum value *σ*_*max*_.

If *p*_*min*_ and *p*_*max*_ are the minimum and maximum values respectively of the PDF under investigation then the function used for computing the standard deviation of the Gaussian kernel based on the PDF *p*(*t*) was defined as:

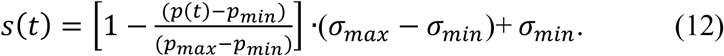

From the first term inside the brackets it is obvious that when the probability *p*(*t*) is low the standard deviation of the Gaussian kernel approaches *σ*_*max*_ while when the probability becomes big, *s*(*t*) approaches *σ*_*min*_.

Based on this function for determining the standard deviation of the blurring Gaussian kernel the probabilistically blurred PDF *p*_*p*_(*t*) was computed as:

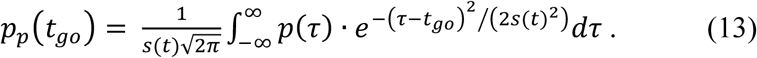

This equation describes that at a given time instance *t*, according to the PDF of event occurrence, the Gaussian uncertainty on the estimation of elapsed time has standard deviation *s*(*t*). The minimum and maximum values of standard deviation *σ*_*min*_ and *σ*_*max*_ are set to *φ* · 0.4 and *φ* · 1.4, as already described earlier. In order to compare this probabilistic blurring hypothesis directly to the initial temporal blurring hypothesis, the value of *φ* was likewise set to 0.21.

Finally, in order to implement the Gaussian blurring of equation (13) at the extrema of ‘go’-times, the definition of the PDF was extended to the left and right of the actual stimulus presentation interval by three standard deviations of the corresponding smoothing Gaussian kernels, similarly to the temporally blurred case described in equation (10), as:

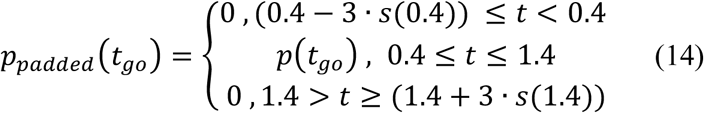

Notice that the extensions depend on the standard deviation function *s*(*t*), which depends on the probability density function.

### Selection of explanatory variables for modeling reaction time as a function of ‘go’ time

The selection of explanatory variables for modeling RT was driven by an expected inverse relation of RT with the PDF and HR of the ‘go’ times. This is based on the rationale that for high relative values of PDF or HR, RT is expected to be small and vice versa. Previous work demonstrated a negative correlation between the ‘subjective’ HR and RT in the order of −0.3 (5). As mentioned earlier, computation of the HR in the brain would require three separate calculations: computation of PDF, its integration for deriving the CDF, and their division for computing HR. The computation of the PDF is the most basic and necessary step in the sequence outlined above and for this reason it was considered here as an alternative factor that can directly affect RT, even before the CDF and HR are computed.

Here two different functions with an inverse character were selected, a ‘mirror’ and a ‘reciprocal’ function. The ‘mirror’ function just reflects a function mirrored around its mean. Here it is used to capture linear anti-correlations between RT and the explanatory variables PDF and HR.

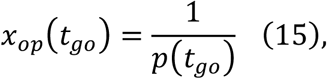

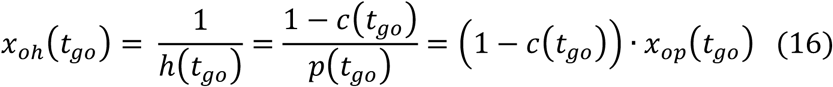

where

*x*_*op*_ : ‘*reciprocal*’*PDF*

*x*_*oh*_ : ‘*reciprocal*’ *hazard ratd of thePDF*

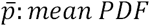, 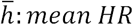

The ‘reciprocal’ function simply takes the reciprocal of a function, e.g. 1 / PDF. This is used to capture a non-linear anti-correlation.

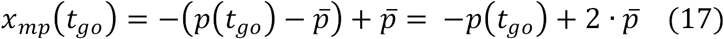

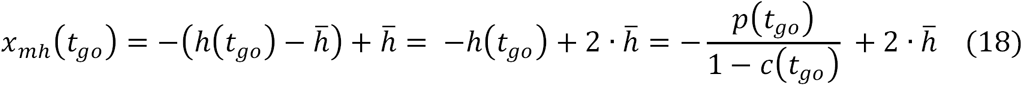

where

*x*_*mp*_: *“mirror” of thePDF*

*x*_*mh*_ : *“mirror” of the hazard rate of the PDF*

Similar variables are defined for the temporally and probabilistically blurred PDF cases, namely *x*_*opt*_, *x*_*oht*_, *x*_*mpt*_, *x*_*mht*_, for temporal blurring and *x*_*opp*_, *x*_*ohp*_ *x*_*mpp*_, *x*_*mhp*_, where for probabilistic blurring where the third letter in the subscript indicates the type of blurring. As it can be seen from equation (16) for any given ‘go’ time *t*_*go*_, the variable *x*_*oh*_(*t*_*go*_) is a scaled version of the variable *x*_*op*_(*t*_*go*_). This scaling depends on the CDF and therefore it is non-linear across ‘go’ times. The same holds for variables *x*_*mp*_(*t*_*go*_) and *x*_*mh*_(*t*_*go*_), as can be seen in equations (17) and (18). As the scaling between these pairs of variables is non-linear (dependent on (1 – *c*(*t*_*go*_)) *and* 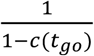 respectively) it is also expected that their relation to RT will not be linearly identical. This also justified their treatment as different variables that are not independent but non-linearly related.

### Modeling RT by ‘mirror’ and ‘reciprocal’ functions with a linear model

The eight ‘blurred’explanatory variables *x*_*opt*_, *x*_*oht*_, *x*_*mpt*_*x*_*mht*_, for temporal blurring and *x*_*opp*_, *x*_*ohp*_ *x*_*mpp*_, *x*_*mhp*_ for probabilistic blurring, were constructed to investigate their linear relation to RT. These variables were derived with *φ* = 0.21 based on previous research (37, 38).

Additionally, the four variables *x*_*op*_, *x*_*oh*_, *x*_*mp*_and *x*_*mh*_ derived directly from the original PDF, CDF and HR, without any Gaussian blurring, were also fit to RT for comparison. A linear model was built for each of these eight explanatory variables. An Ordinary Least Squares (OLS) regression was employed for the computation of the regression coefficients using the MatLab (The MathWorks, Natick MA, USA) *fit* function. Any assumption about the distribution of the residuals of the models was omitted, as we had no evidence that they should follow a Gaussian distribution. Adjusted *R*^2^ was used as a measure of goodness-of-fit for comparing the different models’ relation to RT.

### Comparing temporal and probabilistic blurring

One of the expected differences between the two blurring methodologies was located at the early and late extrema of the ‘go’ time interval. This is because in the case of temporal blurring the smoothing kernel has always much higher variance at the late extremum, as compared to the early, independent of the PDF used and this should be expected to result in greater smearing of RT curves towards the late extremum due to the always greater uncertainty. This should not be the case in probabilistic blurring, where the standard deviation of uncertainty in interval estimation depends on the PDF used. So for example the uncertainty in the early extremum of the exponential PDF should be very similar to that at the late extremum of the flipped-exponential due to the PDF symmetry. This should also result in identical smearing of the RT curves at these two different extrema for these two different PDFs.

In order to investigate if the explanatory variables based on temporally or probabilistically blurred PDFs capture better the behavior of RT curves at different parts of the ‘go’ time range, the ‘go’ time range was divided in 4 equally spaced bins and the goodness-of-fit of all models for the different explanatory variables was computed in each bin. The metric employed for comparing the goodness-of-fit of these models was the sum of squared residuals in each bin. For each distribution (PD_Exp_ and PD_Flip_), each modality (visual, auditory, somatosensory), each of the 4 bins and each of the explanatory variables that was derived based on the blurred PDF, the quotient of the RT model residuals of the temporally blurred case over that of the probabilistically blurred case was computed. This procedure indicated which blurring method provided the model that fits the RT of a specific bin better.

