## Supplementary Material for "The Anticipation of Events in Time"

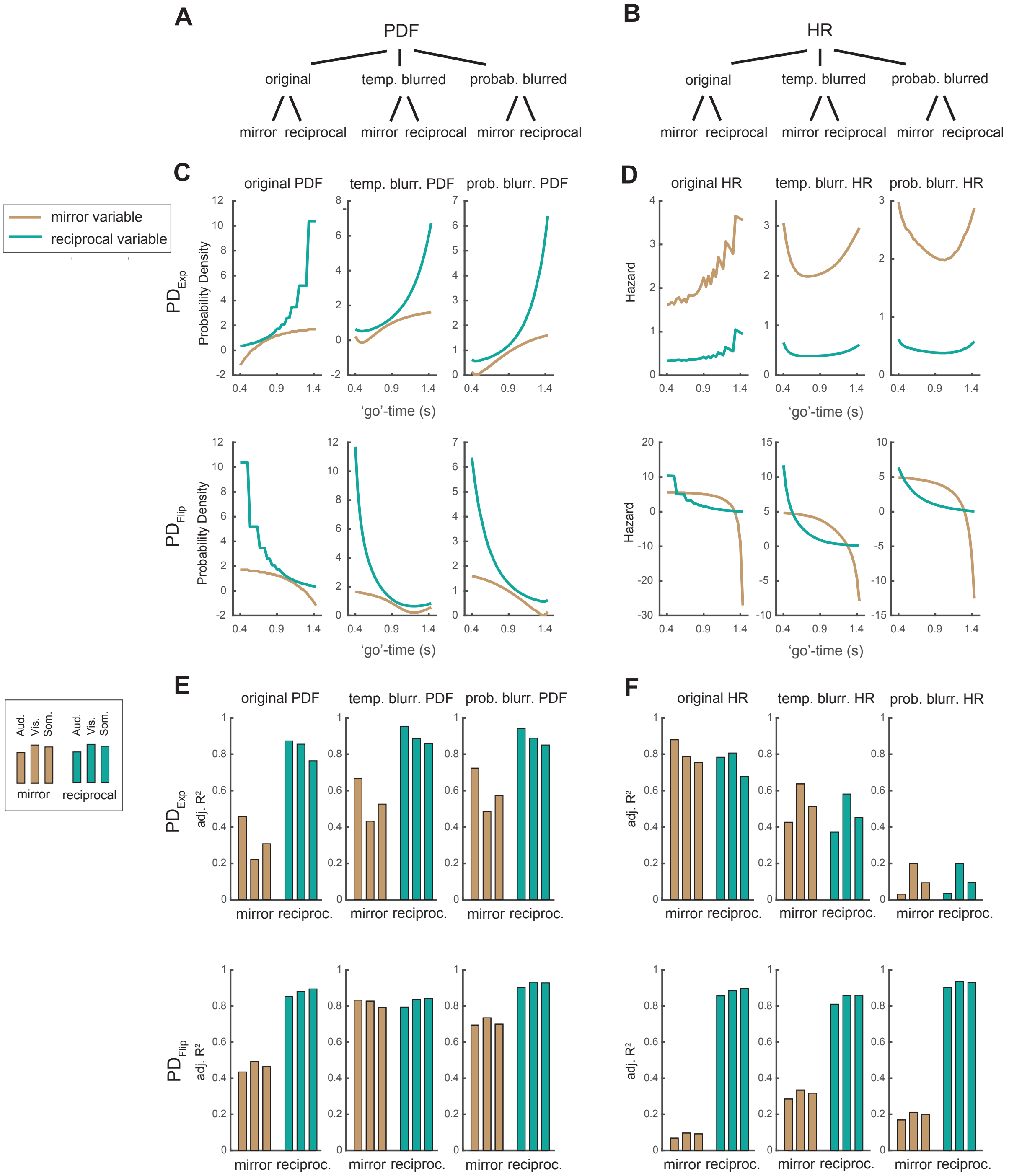


**Fig. S1**. Models of reaction time with respect to probability over time. (A) and (B) Schematic illustrating the relationship between a presented PDF or its HR and the derived explanatory variables used to model RT. (C) Exponential and flipped exponential PDFs in original, temporally and probabilistically blurred versions. The original presented 'go' time functions and their blurred versions are shown to highlight the effect of the convolution with a Gaussian distribution whose standard deviation scales either with 'go' time (temporal blurring) or it scales according to the PDF of event occurrence (probabilistic blurring). In case of the PDFs the blurring changes the monotonically decreasing ($PD_{Exp}$) and increasing ($PD_{Flip}$) shapes to biphasic curves. (D) In the HRs, the blurring had a stronger effect in the $PD_{Exp}$ condition than in the $PD_{Flip}$ condition. Based on the assumption that RT should be smallest where probability – or hazard – is highest, an inverse relationship between RT and model was considered. Before fitting the models to the data, all explanatory variables were either mirrored around their mean, which is a linear transformation, or they were transformed into their reciprocal versions, which is a nonlinear transformation. (E) and (F) Adjusted $R^2$ of the model fits. Each of the explanatory variables was fitted to RT of the respective condition ($PD_{Exp}$ or $PD_{Flip}$) in all sensory conditions.

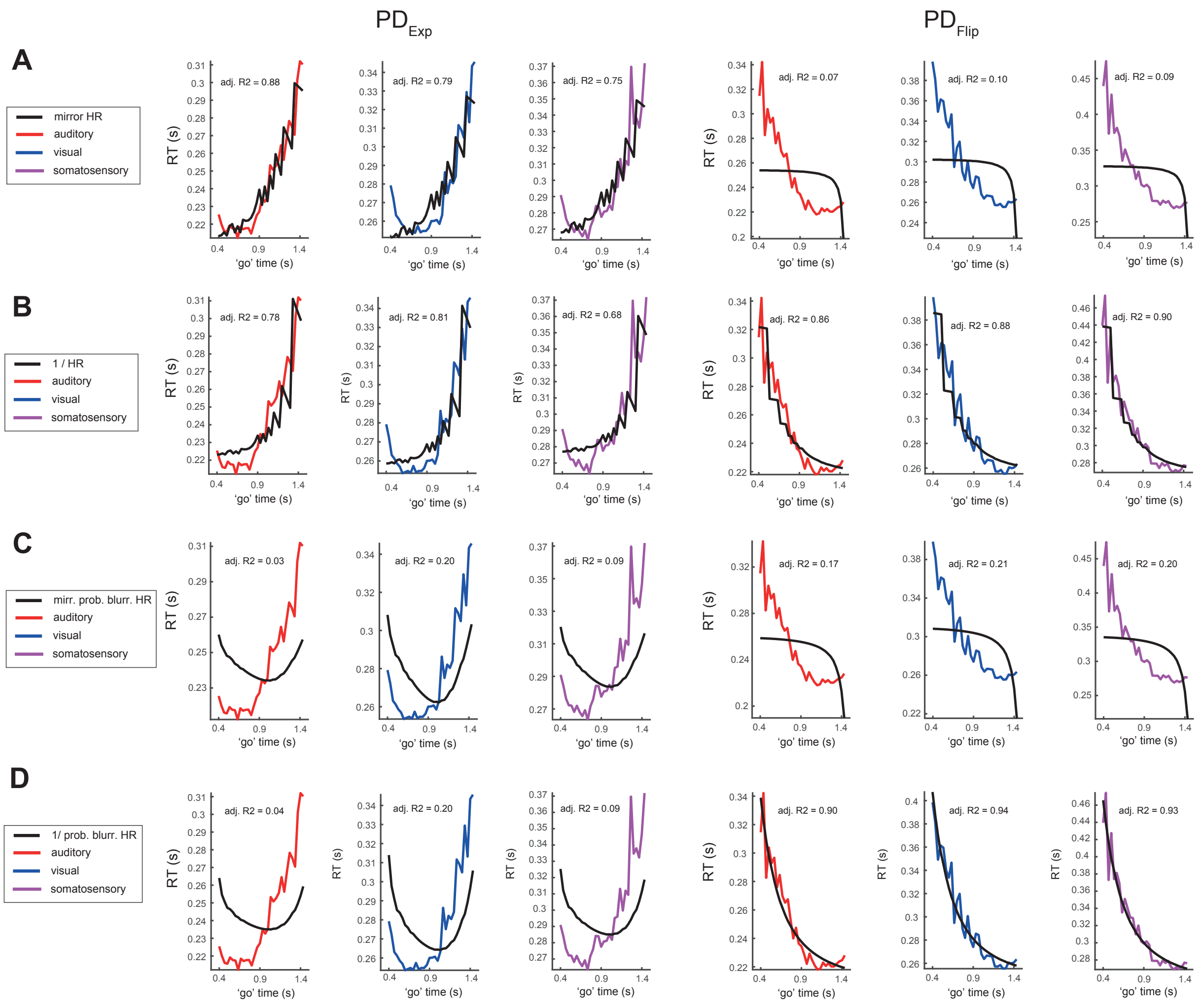


**Fig. S2**. Models of RT based on the hazard rate (HR) of 'go' times. (A) The normal HR does not capture the data well in the $PD_{Flip}$ condition. (B) The reciprocal of the HR fits the data better than the mirror one. In the $PD_{Exp}$ condition the model does not capture RTs at shorter 'go' times. Although in the $PD_{Flip}$ condition the model fits the data better and thus gives a relatively high adjusted $R^2$, the stepped character of the explanatory variable is not reflected in the data. (C) The predictions of the model does not match the data in either the $PD_{Exp}$ condition, or the $PD_{Flip}$ condition. (D) The probabilistically-blurred HR does not capture the pattern of RT in the $PD_{Exp}$ condition. Although in the $PD_{Flip}$ condition the model clearly fits the data better and thus gives a high adjusted $R^2$, a convincing model needs to capture the RT data in both probabilistic conditions.

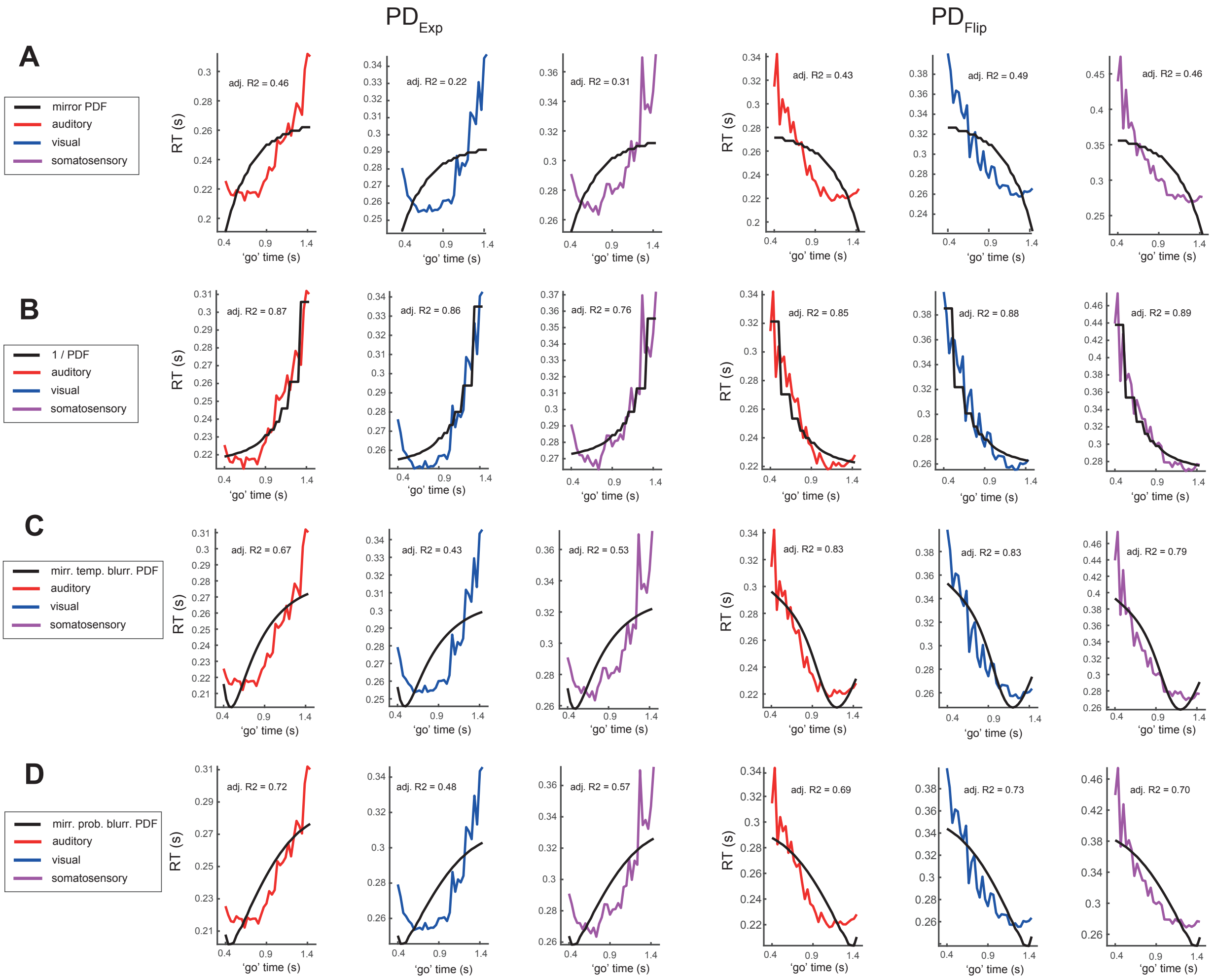


**Fig. S3**. Models of RT based on the probability density function (PDF) of 'go' times. (A) The mirrored original PDF (linearly transformed) does not fit the data in either the $PD_{Exp}$ condition or in the $PD_{Flip}$ condition. (B) The original PDF captures the behavior of the RT data well in both the $PD_{Exp}$ condition and the $PD_{Flip}$ condition, as evidenced by the high adjusted $R^2$. However, the original PDF contains step-discontinuities not evident in the data. (C) The mirrored, temporally-blurred PDF does not fit the data in either the $PD_{Exp}$ condition or in the $PD_{Flip}$ condition. (D) The mirrored probabilistically-blurred PDF does not fit the data well in either the $PD_{Exp}$ condition or in the $PD_{Flip}$ condition.

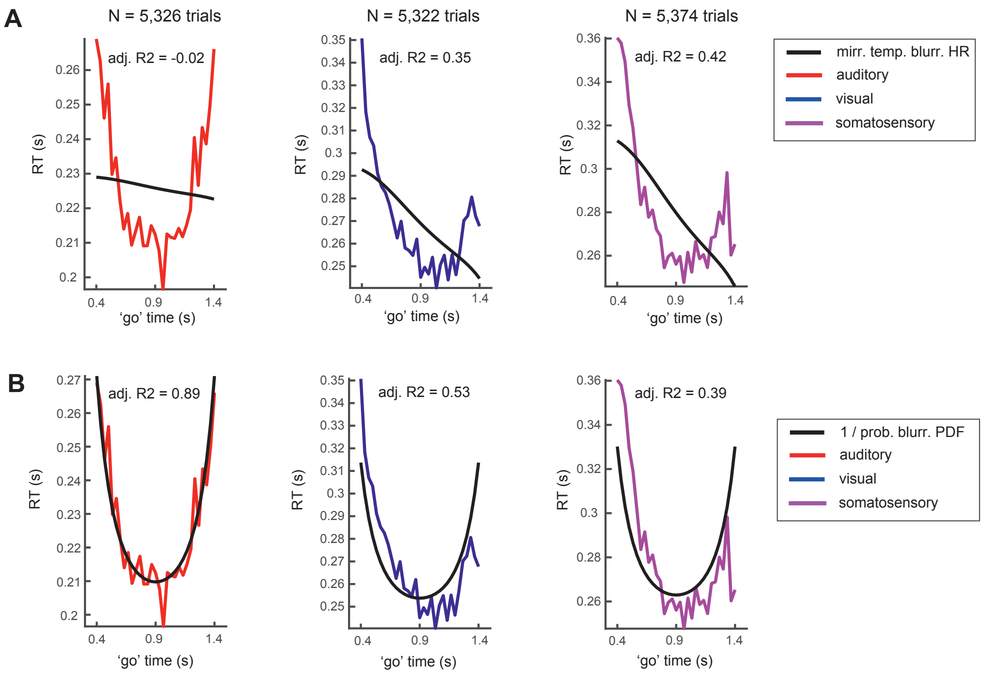


**Fig. S4**. In Gaussian condition, models based on probabilistically-blurred, reciprocal PDF capture the reaction time data better than the temporally-blurred, mirrored hazard rate. Auditory (red), visual (blue), and somatosensory (violet) reaction time data from 18 participants who performed the 'set' - 'go' task with a Gaussian distribution of 'go' times (see Methods). (A) The *temporally-blurred, mirrored hazard rate* cannot account for the reaction time modulation in any of the three modalities. (B) The *probabilistically-blurred, reciprocal PDF* fits reaction times better in all three modalities, especially in the auditory condition. Although in visual and and somatosensory conditions the model fits the data less accurately, it still captures the biphasic reaction time modulation, a feature absent in the mirrored HR case (A).

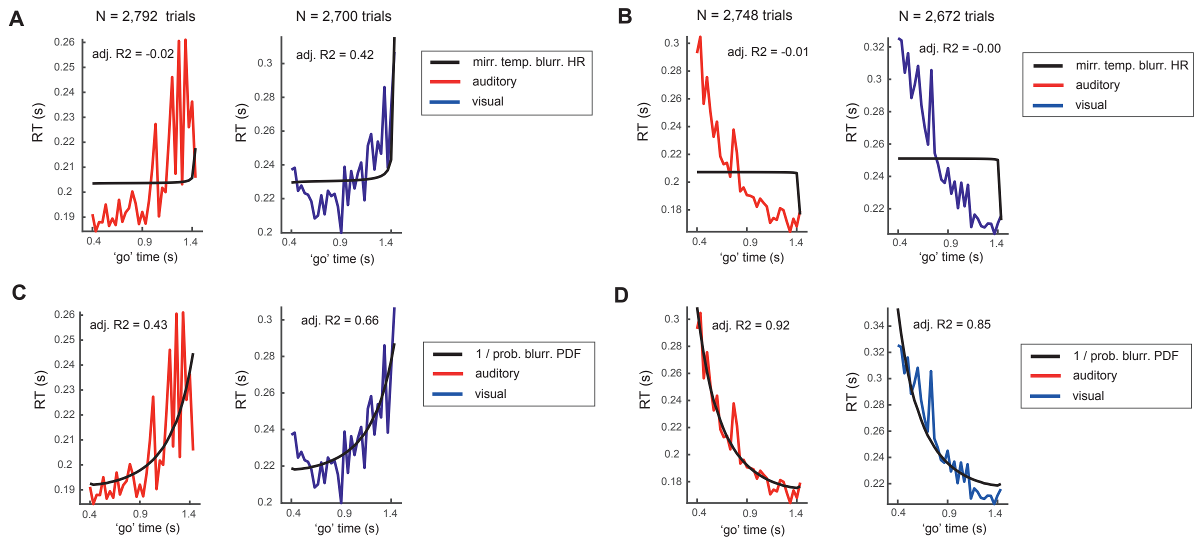


**Fig. S5**. In deterministic setting, models based on hazard rate fail to capture RT data while models based on PDF fit the data well. (A to D) Auditory (red) and visual (blue) reaction time data from 18 participants who performed the 'set' - 'go' task without catch trials. In this *deterministic setting*, where there in no uncertainty about 'go' cue occurrence, exponential ($PD_{Exp}$) and flipped-exponential ($PD_{Flip}$) distributions of 'go' time were presented in audition and in vision. (A and B) In neither $PD_{Exp}$ (A), nor $PD_{Flip}$ (B) conditions does the *temporally-blurred, mirrored hazard* rate fit the data. (C and D) The *probabilistically-blurred, reciprocal PDF* captures reaction times adequately in all conditions.

| | *Probability distribution* | | *Sensory modality* | | *Day* | |
|---|---|---|---|---|---|---|
| | $F_{(1,287)}$ | $P$ | $F_{(2,287)}$ | $P$ | $F_{(1,287)}$ | $P$ |
| median RT | 0.25 | 0.64 | 41.70 | $1.3*10^{-16}$ | 22.10 | $4.1*10^{-6}$ |
| $IQR_{RT}$ | 8.40 | 0.0041 | 14.50 | $1.0*10^{-6}$ | 29.30 | $1.3*10^{-7}$ |
| $\mu$ | 0.42 | 0.52 | 63.8 | $1.3*10^{-23}$ | 11.80 | 0.0007 |
| $\sigma$ | 0.93 | 0.34 | 12.40 | $7.1*10^{-6}$ | 3.99 | 0.049 |
| $\tau$ | 6.51 | 0.011 | 10.82 | $3.0*10^{-5}$ | 30.64 | $7.1*10^{-8}$ |

**Table S1**. Three-way analysis of variance (ANOVA) on factors probability distribution, sensory modality and day. All three-way and two-way interaction were n.s. and were therefore removed from all above models.

| | Auditory | $t_{(23)}$ | $P$ | Visual | $t_{(23)}$ | $P$ | Somato. | $t_{(23)}$ | $P$ |
|---|---|---|---|---|---|---|---|---|---|
| $\Delta$ $IQR_{RT}$ | -10.3 (19.4) | -2.62 | 0.0154 | -11.7 (20.2) | -2.85 | 0.0091 | -11.4 (12.7) | -4.39 | 0.00022 |
| $\Delta$ $\tau$ | -9.7 (18.5) | -2.58 | 0.0166 | -9.5 (17.5) | -2.67 | 0.0136 | -8.2 (13.7) | -2.92 | 0.0077 |

**Table S2**. *t* tests on significant factors (three-way ANOVA, planned contrasts). Differences in variables between probabilistic conditions ($PD_{Exp}$ - $PD_{Flip}$). All variables are in units of ms, all variances (in parentheses) are standard deviations.

| | Aud. - Vis. | $t_{(23)}$ | $P$ | Aud. - Som. | $t_{(23)}$ | $P$ | Vis. - Som. | $t_{(23)}$ | $P$ |
|---|---|---|---|---|---|---|---|---|---|
| Δ median RT | -43.2 (24.5) | -8.7 | $1.1 * 10^{-8}$ | -55.0 (29.2) | -9.2 | $3.4 * 10^{-9}$ | -11.7 (35.6) | -1.6 | 0.12 |
| Δ $IQR_{RT}$ | 7.5 (20.5) | 1.78 | 0.089 | -14.1 (19.4) | -3.56 | 0.0019 | -21.5 (18.2) | 5.80 | $6.6*10^{-6}$ |
| Δ $\mu$ | -47.7 (20.1) | -11.6 | $4.1*10^{-11}$ | -44.9 (24.7) | -8.9 | $6.4*10^{-9}$ | 2.8 (28.8) | 0.49 | 0.64 |
| Δ $\sigma$ | 1.3 (6.7) | -0.98 | 0.34 | -6.7 (10.3) | -3.17 | 0.0043 | -5.3 (9.8) | -2.66 | 0.014 |
| Δ $\tau$ | 7.1 (22.0) | 1.58 | 0.13 | -12.5 (20.3) | -3.01 | 0.006 | -19.5 (15.9) | -6.01 | $3.9 *10^{-6}$ |

**Table S3**. *t* tests on significant factors (three-way ANOVA, planned contrasts). Differences in variables across sensory modalities within $PD_{Exp.}$ All variables are in units of ms, all variances (in parentheses) are standard deviations.

| | Aud. - Vis. | $t_{(23)}$ | *P* | Aud. - Som. | $t_{(23)}$ | *P* | Vis. - Som. | $t_{(23)}$ | *P* |
|---|---|---|---|---|---|---|---|---|---|
| Δ median RT | -41.7 (29.0) | -7.0 | $3.7 * 10^{-7}$ | -54.7 (38.4) | -7.0 | $4.1 * 10^{-7}$ | -13.1 (47.9) | -1.3 | 0.20 |
| Δ $IQR_{RT}$ | 6.0 (35.5) | 0.84 | 0.41 | -15.1 (32.0) | -2.31 | 0.0302 | -21.2 (33.1) | -3.14 | 0.0047 |
| Δ $\mu$ | -45.9 (21.3) | -10.5 | $2.8*10^{-10}$ | -43.9 (29.1) | -7.4 | $1.7*10^{-7}$ | 2.1 (34.8) | 0.29 | 0.78 |
| Δ $\sigma$ | -1.1 (6.7) | -0.82 | 0.42 | -9.3 (11.8) | -3.87 | 0.0008 | -8.2 (11.8) | -3.42 | 0.0024 |
| Δ $\tau$ | 7.3 (33.3) | 1.07 | 0.30 | -10.9 (31.2) | -1.72 | 0.10 | -18.2 (30.0) | -2.98 | 0.0068 |

**Table S4**. *t* tests on significant factors (three-way ANOVA, planned contrasts). Differences in variables across sensory modalities within $PD_{Flip.}$ All variables are in units of ms, all variances (in parentheses) are standard deviations.

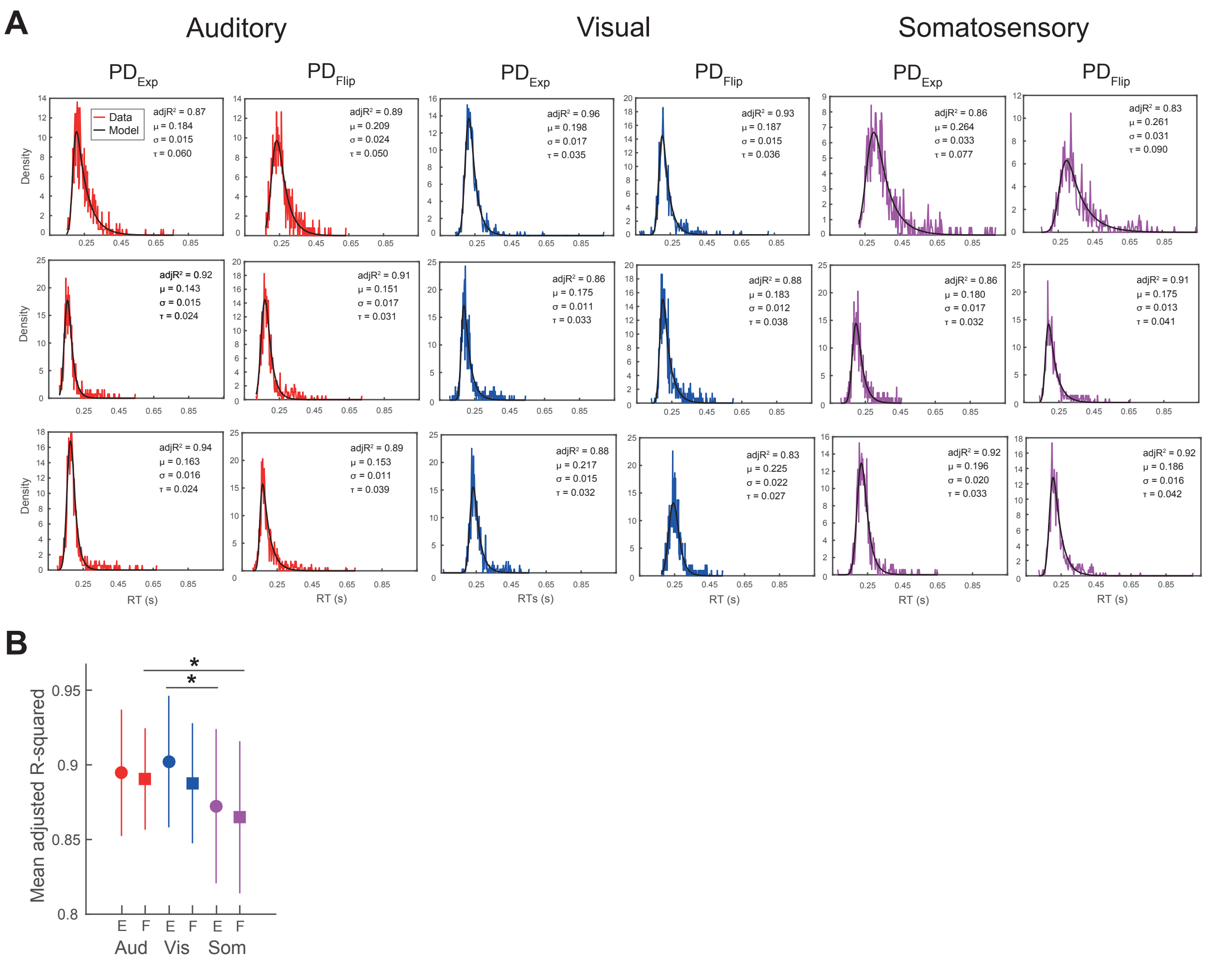


**Fig. S6**. (A) Exemplary exponential-Gaussian fits to RT. Each row contains data for a single subject. In the three sensory modalities and in the two probabilistic conditions, the ex-Gaussian model captured the data well. (B) Goodness-of-fit of ex-Gaussian model. RT data were well-fit by the ex-Gaussian model in all sensory modalities and in both probabilistic conditions as shown by the overall high $R^2$ (planned contrasts, * $P < 0.05$, $t$ tests (two tailed), error bars, ± SD).

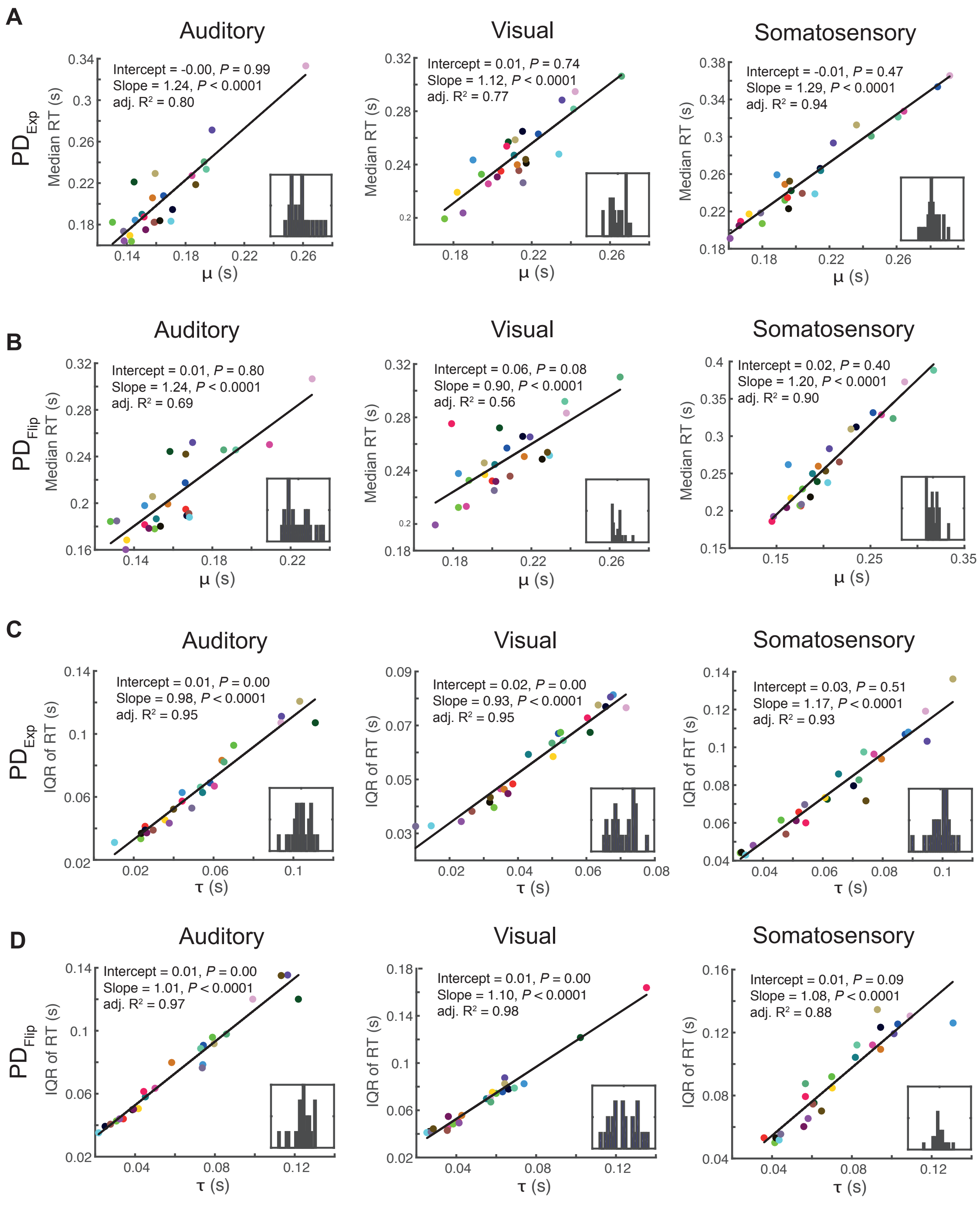


**Fig. S7**. Ex-Gaussian fit parameters $\mu$ and $\tau$ capture average RT and variance of RT. (A and B) In both $PD_{Exp}$ and $PD_{Flip}$ conditions, the relationship between Gaussian $\mu$ and median RT is captured adequately by a linear model (black fit line). (C and D) Similarly, in all conditions, $\tau$ and $IQR_{RT}$ are linearly related (black fit line). Each dot represents a single subject. Inset graphs show residuals of linear fit.

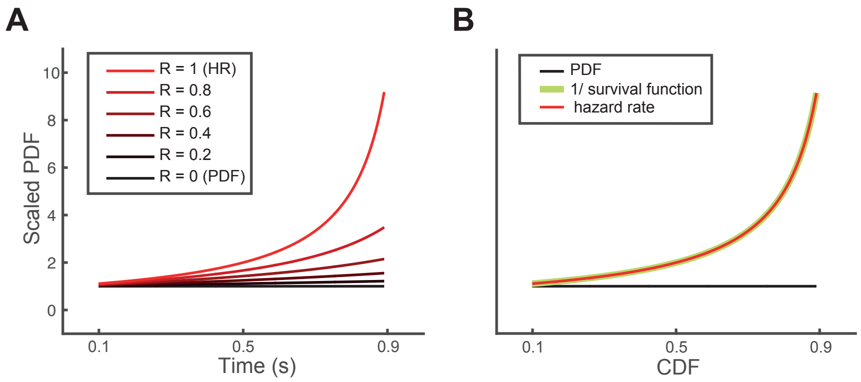


**Fig. S8**. Hypothesized effect of reward expectation on probability density and influence of survival function on hazard rate. (A) Influence of R, a factor hypothesized to reflect reward expectancy, on probability density. The parameter described by the formula PDF/(1-R*CDF), is termed here the *scaled PDF* in which the term 1/(1-R*CDF) is the scaling factor. In the case of no expected reward, R approaches 0, the scaling factor approaches a constant value across time, and the *scaled PDF* approaches the PDF itself. In the case of high reward expectation, R approaches 1 and the *scaled PDF* approaches the HR, as HR = PDF/1-CDF. (B) Functions of a uniform PDF that spans from zero to one. In the formula for the HR, HR = (1/survival function) * PDF, the scaling term 1/survival function monotonically increases, with the highest gradient occuring when CDF approaches one at the right extremum of any distribution. These high scaling values drive the hazard rate to also sharply increase (red line). This highlights the dominant role of  the term 1/survival rate on the values of HR towards the right extremum of PDFs. Especially in the absence of catch trials, where the CDF approaches one asymptotically, the factor 1/survival rate takes extremely high values near the end of the distribution (survival rate = 1-CDF), thus outweighting the impact of the PDF (at least for the majority of PDFs). For plotting, the functions have been truncated to avoid extreme values at the right extremum. The y-axis accomodates probability density, cumulative probability density and hazard rate, therefore no units are specified.

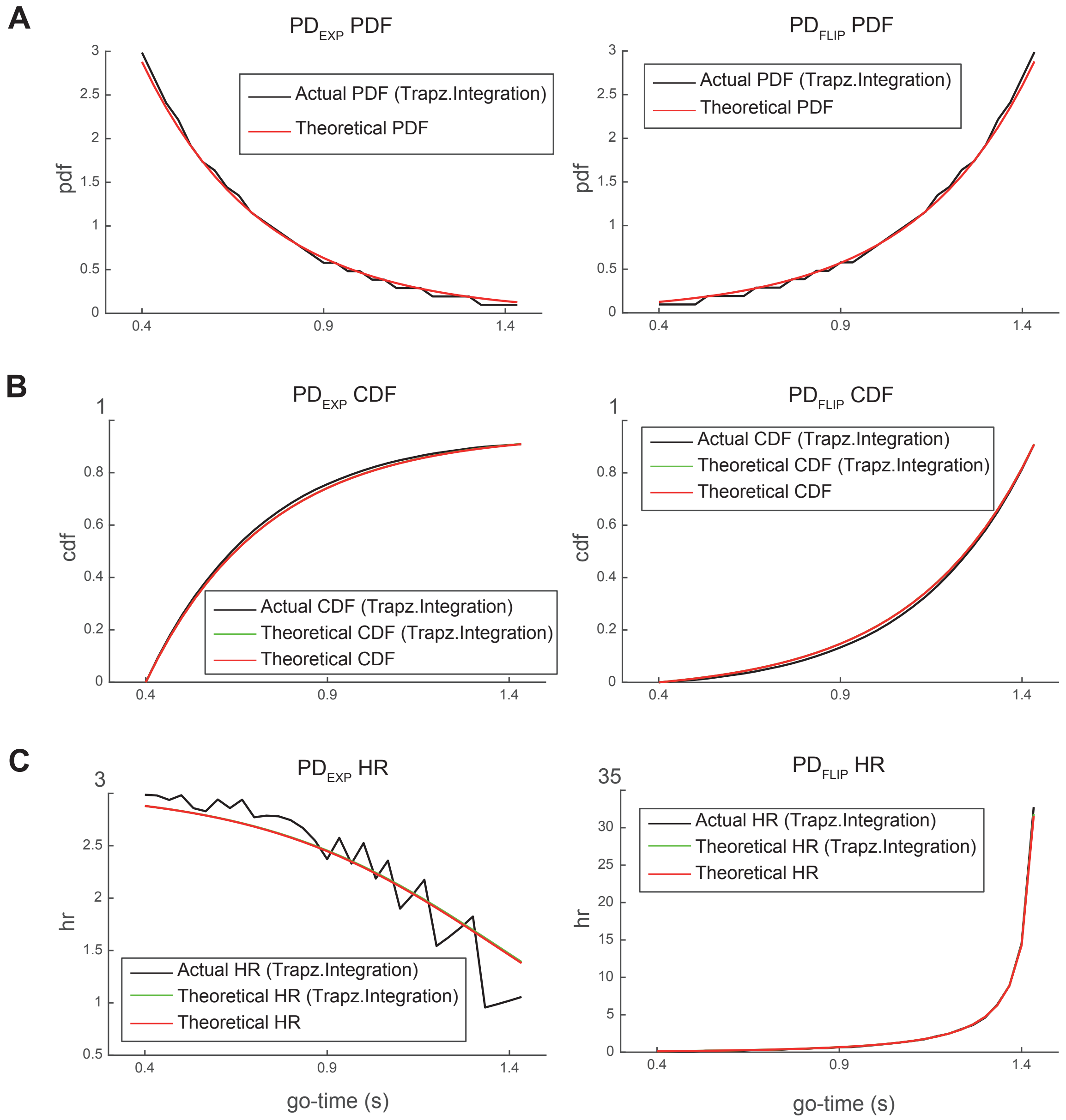


**Fig. S9**. Comparison of theoretical PDF, CDF and hazard rate (HR) with the discretized, versions presented in the experiment. The discretized versions of the CDF were derived by trapezoidal integration of the discretized PDF. (A) The discretized versions of the PDF approximated closely the theoretical ones (notice the step-like shape). (B) The same was true for the CDF (since the discretized CDF is the integral of the discretized PDF it is not itself a step function but rather a piecewise linear function). (B) The jigsaw-pattern in the HR of the $PD_{Exp}$ condition is a consequence of the discretization of the PDF while the CDF remains continuous. Regardless of these jigsaw-pattern the discretized HR curve follows the theoretical one.
